# *hsp-16.2* chaperone biomarkers track physiological states of proteome dosage

**DOI:** 10.1101/431643

**Authors:** Nikolay Burnaevskiy, Bryan Sands, Soo Yun, Patricia M Tedesco, Thomas E Johnson, Matt Kaeberlein, Roger Brent, Alexander Mendenhall

**Affiliations:** Department of Pathology, University of Washington, Seattle, Washington; Department of Integrative Physiology, University of Colorado, Boulder, Colorado; Division of Basic Sciences, Fred Hutchinson Cancer Research Center, Seattle, Washington

## Abstract

Phenotypic expression of many traits varies among isogenic individuals in homogeneous environments. Intrinsic variation in the protein chaperone system affects a wide variety of traits in diverse biological systems. In *C. elegans*, expression of *hsp-16.2* chaperone biomarkers predicts the penetrance of mutations and lifespan after heat shock. But the physiological mechanisms by which cells express different amounts of the biomarker were unknown. Here, we used an *in vivo* microscopy approach to dissect the mechanisms of cell-to-cell variation in *hsp-16.2* biomarker expression, focusing on the intestines, which generate most signal. We found both intrinsic noise and signaling noise are low. The major axis of cell-to-cell variation in gene expression is composed of general differences in protein dosage. Thus, *hsp-16.2* biomarkers reveal states of high or low effective dosages for many genes. It is possible that natural variation in protein dosage or chaperone activity may account for missing heritability of some traits.

## INTRODUCTION

Genes and environments do not explain all the differences in heritable traits. Experiments with isogenic model systems kept in homogeneous environments have clearly demonstrated this^1^. Variation in traits among isogenic animals was perhaps first noticed in 1925 in *Drosophila funeberis* by Romaschoff when he noticed that not all animals in a “pure bloodline” (inbred strain) exhibited the mutant phenotype for *Abdomen abnormalis*; this is called incomplete penetrance. The severity of the phenotype was different as well, a phenomenon that came to be termed expressivity^2^. Also in 1925, Timofeeff-Resskovsky noticed that the penetrance and expressivity of another mutation, *Radius incompletes*, was altered in different genetic backgrounds. Thus, the action of genes could affect the penetrance and expressivity of discreet traits conferred by other genes^3^. Complex traits such as lifespan also vary among isogenic individuals, both in model systems^4^ and in human monozygotic twins^5^. The mechanisms underlying this variation remain poorly understood.

Experimental and natural variation of genes in the protein chaperone system alters the manifestation of complex and discreet traits. In the late 1990s and 2000s scientists found that perturbing the chaperone system had broad effects on variation in traits, including the penetrance of some mutations. Scientists used geldanamycin to perturb the N-terminus of HSP90, a chaperone with important specific clients, and a master regulator of the heat shock response^6^. They found that the general variation in traits is affected in different genetic backgrounds of *Drosophila melanogaster*^7^ and *Arabidopsis thalania*^8^. Another group found chaperone-altering geldanamycin treatment alters penetrance and expressivity of specific mutations in the vertebrate *Danio rerio*^9^. In *Caenorhabditis elegans*, which is immune to the effects of geldanamycin, differences in levels of *hsp-90* reporter gene expression levels predicted differences in the penetrance of loss of function mutations^10,11^ (but not null mutations; this is important to consider for interpretation).

In previous work, we explored the consequences of differences in expression of another chaperone on expression of genetic traits. In *C. elegans*, the *hsp-16.2* reporter gene is expressed only after heat shock. We found that adult animals that make more of the *hsp-16.2* reporter gene have differences in complex traits – lifespan and lethal thermal stress tolerance in *C. elegans*^12,13^. Using the another construct of the *hsp-16.2* reporter in *C. elegans* (the same promoter fused to fluorescent protein, inserted elsewhere in the genome), another group found that increased *hsp-16.2* reporter expression was associated with differences in the penetrance of a number of hypomorphic point mutations in distinct types of genes^10^. For the most part, these *hsp-16.2* and *hsp-90* reporter gene biomarkers correlated with the penetrance of distinct mutations, but both correlated with penetrance of at least one mutation, *lin-31*^10^, indicating that there are both distinct and overlapping fractions of the proteome affected by these chaperones. As expected, but important to note, both *hsp-90*^11^ and *hsp-16.2*^12^ reporters properly correlate with the expression levels of the chaperones on which they are reporting.

We previously showed that differences in expression levels of *hsp-16.2* lifespan/penetrance biomarkers in adult animals were likely due to differences in transcription; notably, this did not include *hsp-90*. We also identified genes^14^ and environmental variables^15^ that affected the amount of interindividual variation in the *hsp-16.2* reporter. Yet, we did not know how the cells of animals came to express more or less of this lifespan/penetrance biomarker. Therefore, we set out to dissect the mechanisms of cell-to-cell variation in gene expression to understand how differences in the expression of *hsp-16.2* lifespan/penetrance biomarker arise. We focused on gene expression in the intestine cells of adult animals because that is the tissue that makes the most signal for *hsp-16.2* reporters^13,16,17^, and because it is the point in life we used to predict lifespan and thermotolerance^12,13^, and because we had developed technical methods for *in vivo* reproducible quantification of gene expression in single intestine cells^16^.

Here, we used an experimental design and analytical framework derived from yeast^18^, which allowed us to discern between three ideas for how differences in *hsp-16.2* reporter expression might arise. The first hypothesis was that differences in the *hsp-16.2* lifespan biomarker might arise from differences in intrinsic noise in gene expression. Previous work in yeast^19^ and humans^20^ showed that individual cells may only express much less of, or only one, of their two distinct copies of each allele. Therefore, animals might express more or less of a gene by expressing different amounts of each copy – anywhere from full expression of both alleles to no expression of either allele.

Our second hypothesis was that differences in the *hsp-16.2* reporter and associated chaperones might be due to differences specific to the signaling pathway that activated chaperone expression. That is, we hypothesized we would see relatively high covariation for expression of the *hsp-16.2* reporter gene and other chaperone reporters like *hsp-90*, but not covariation with non-heat shock pathway reporters like a yolk-protein reporter.

Finally, our third hypothesis was that elevated chaperone levels would increase general protein dosage, by affecting expression capacity or protein turnover. Chaperones have many clients and might affect folding, maintenance and turnover of these proteins^6^, thereby altering their effective dosage. This idea was appealing, since, work from the Kim Lab using *C. elegans* found that several distinct non-chaperone reporter genes could predict lifespan, and that these distinct reporters were highly correlated^21^. Moreover, work by us in *S. cerevisiae* had shown that these general effects on protein dosage^18^ are important contributors to extrinsic noise in gene expression in *E. coli*^22^ and *S. cerevisiae*^19^. A detailed description of this analytical framework is given in Supplementary Materials Section 2. Cartoon images of what cell autonomous and cell nonautonomous differences in each of the three aforementioned categories (intrinsic noise, signaling noise, and general protein dosage) are shown as Supplementary Figs. S1-S3.

Below, we detail results showing that, for *hsp-16.2* reporter expression in the adult worm intestine, two components of cell-to-cell variation are minimal. The other component, differences in protein dosage, accounts for the majority of variation in gene expression in intestine cells. We provide experimental evidence that shows how differences in this component may arise after heat shock in the context of a working model integrating data from this and other reports, and suggest how these differences might account for observed effects on expressivity and penetrance of different alleles.

## RESULTS

### Adaptation of an Analytical Framework and Experimental Design

Here we adapted an approach we used in yeast^18^, wherein we compared the outputs of two differently colored (different fluorescent proteins) versions of the same reporter gene expressed from two identical loci on homologous chromosomes (Type I experiments) or the outputs of two different reporter genes (Type II experiments), shown in Fig. 1. Measuring the pairs of reporter genes in these two different kinds of experiments allows us to quantify the amount of cell-to-cell variation in gene expression attributable to three distinct, experimentally tractable bins, each with distinct underlying molecular causes. Fig. 1 shows how we can use this reporter gene measurement scheme to quantify differences in intrinsic noise^22,23^ (η^2^(γ)), signaling through different pathways^18^ (η^2^(P)), or general protein expression capacity^18^, a measure of general protein abundance (η^2^(G)). See Supplementary Materials Section 2: Analytical Framework and Supplementary Figs. S1-S3 for additional explanation and cartoon depictions of different kinds of variation in gene expression.

**Figure 1.**
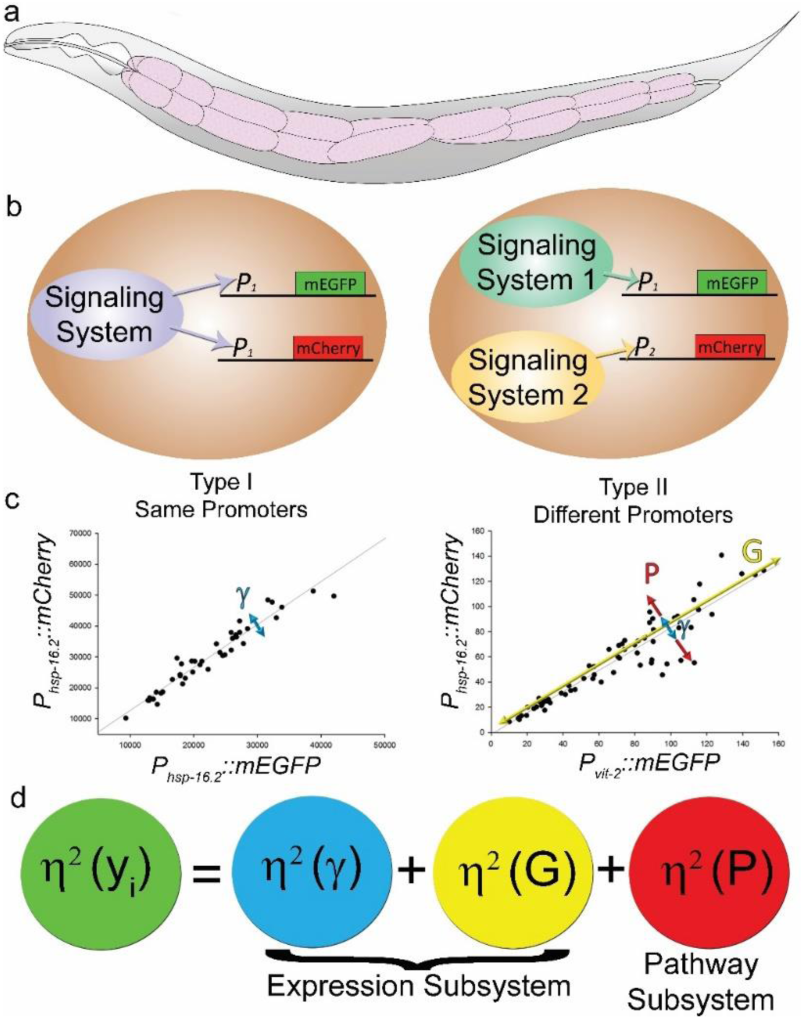
Experimental design and analytical framework. **a.** Schematic view of the *C. elegans* body. Intestine is colored pink. The twenty cells of the intestine are organized into nine segments, called rings. **b.** Overview of reporter gene measurement scheme. In the left panel, two differently colored copies of the same gene respond to the same signaling system (Type I experiments), or, in the right panel two differently colored reporters controlled by two different promoters respond to two distinct signaling systems (Type II experiments)**. c.** Data plotting schematic. We imaged intestine cells in rings one through four (the cells that fit in a single field of view with a 40x objective) in animals on day two of adulthood, at the point when lifespan can be predicted from *P*_*hsp-16.2*_::*gfp* expression levels. We then plotted expression levels of reporter pairs expressed in the intestine cells, grouped by cell type; the same cell types (e.g., cells in ring three; int3 cells) from different animals are plotted. Left scatterplot shows typical results of type I experiment. Since both genes receive the signals from the same upstream regulators and share the same downstream expression machinery, uncorrelated variation of expression results only from stochastic noise of transcription/translation or variable allele access η^2^(γ). Right scatterplot shows typical results of type II experiment. In the case of two different genes, uncorrelated variation of expression can result from stochastic noise of transcription/translation or variable allele access η^2^(γ) and variation in activation of particular signaling pathways η^2^(P). *C*orrelated variation results from variation in shared gene expression machinery η^2^(G). **d.** A simplified version of the analytical framework derived from Colman-Lerner et al 2005 is shown; it depicts the three experimentally tractable bins into which we can attribute cell-to-cell variation in gene expression. Each bin has different underlying molecular causes. A complete, mathematically detailed description of the analytical framework is shown in Supplementary Materials Section 2.

### Type I Experiments Quantify Intrinsic Noise (η^2^(γ))

Type I experiments allow us to measure how much cell-to-cell variation there is in apparently stochastic to-promoter or to-transcript binding events, or allele access. Type I experiments reveal how much cell-to-cell variation there is in allele expression/access as uncorrelated variation, often referred to as intrinsic noise (η^2^(γ)); Fig. 1 shows an example of a scatterplot for the same cell type (same lineage/fate) measured from many different animals from one of these experiments. If there were a lot of uncorrelated variation (a cloud like appearance of points) it would mean that there were significant cell-to-cell differences in to-promoter binding, to-transcript binding, or allele access. If there was little uncorrelated variation (alignment of points along the major axis; Fig. 1), it would mean that most cells would have had access to both alleles. It would also mean that there were few differences attributable to probabilistic differences in biochemical binding events (to-promoter/to-transcript) that tend to dominate when there are small numbers of molecules involved^23^; we would say that intrinsic noise was low.

### Intrinsic Noise in Gene Expression is Relatively Constrained in *C. elegans* Intestine Cells

In Type I experiments for *hsp-16.2*, and for all the other reporter genes we measured, intrinsic noise was low. The expression level of one allele accounted for 90% or more of the expression of the other allele in animals. Fig. 2 shows the average intrinsic noise levels for a few genes we measured. Intrinsic noise was a minor component of the cell-to-cell variation, and relatively constrained compared to yeast^19^. We show individual Type I scatterplots and bar graphs organized by cell type and experiment in Supplementary Figs. S4&S5. A few relatively deviant cells can be seen off the trendline in Fig. S4, which shows the same cell measured from many different animals across multiple experiments, demonstrating that this experimental system is capable of detecting intrinsic noise in gene expression at the protein level. However, large differences in allele bias simply do not occur at a high frequency in the intestine cells, also shown by Supplementary Fig. S4.

**Figure 2.**
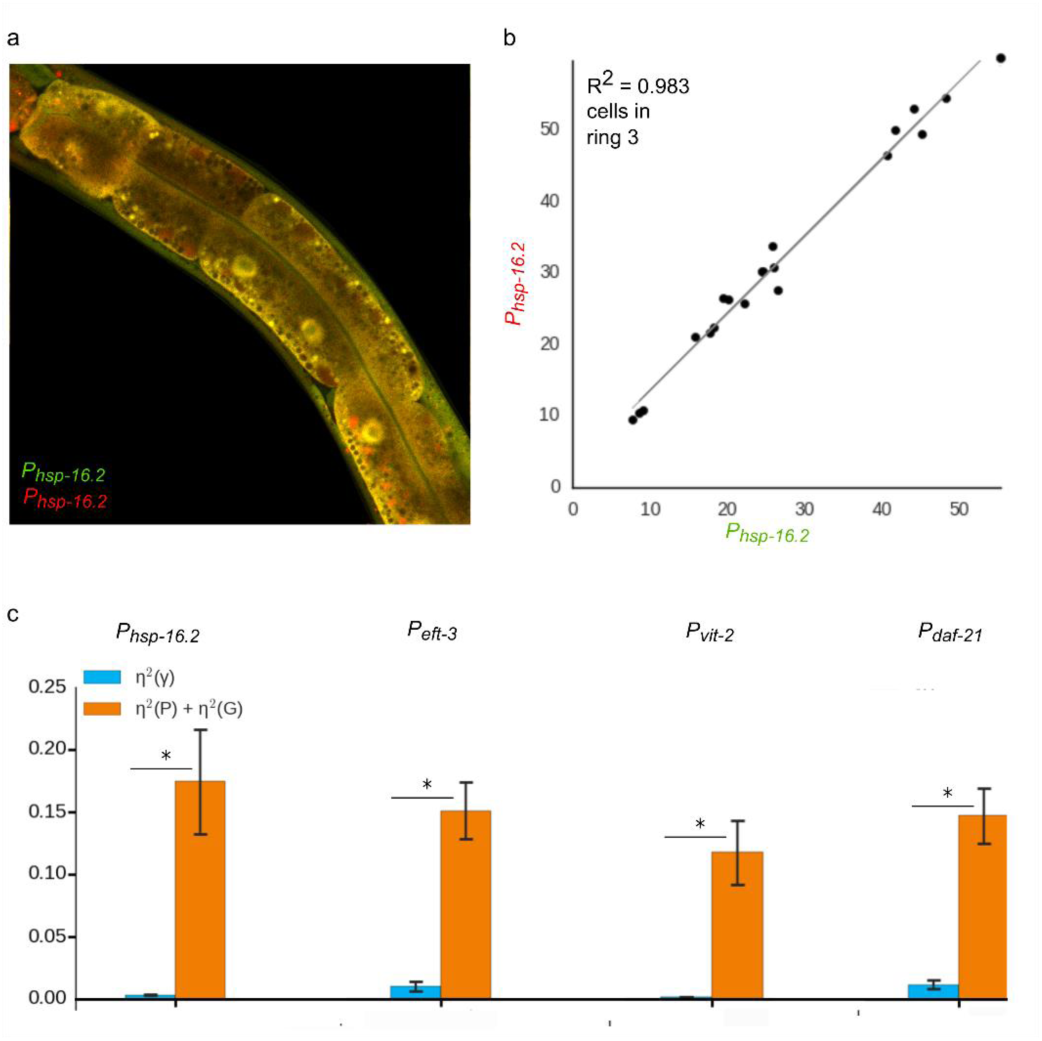
Type I experiments reveal low intrinsic noise. **a.** An example of the animal producing fluorescent reporters (mEGFP and mCherry) from two identical promoters (*P*_*hsp-16.2*_). We did not observe patchwork of green and red cells. Cells were nearly uniformly yellow, indicating low intrinsic noise. **b.** Scatterplot of *P*_*hsp-16.2*_::GFP and *P*_*hsp-16.2*_::mCherry expression in cells in ring 3 of the intestine from ten animals examined in one experiment. **c.** Average correlated and uncorrelated variations for *P*_*hsp-16.2*_, *P*_*eft-3*_, *P*_*vit-2*_ and *P*_*daf-21*_ in the intestine cells in rings 1-4. In type I experiment uncorrelated variation arises from stochastic noise of transcription/translation or variable allele access – η^2^(γ); correlated variation is a combined result of variation in gene expression capacity η^2^(G) and variation in pathway activation η^2^(P). * indicates statistical significance *P* < 0.05 by two-tailed t-test; error bars are S.E.M.. For each pair of alleles, data are from at least three independent experiments measuring eight intestine cells in at least ten animals per experiment.

### Type II Experiments Quantify Signaling Noise (η^2^(*P*)) and Differences in Protein Expression Capacity (η^2^(G))

Type II experiments compare the expression levels from two distinct genes to tell us about cell-to-cell differences in signaling through different pathways (η^2^(*P*)). In the scatterplot from a Type II experiment shown in Fig. 1, the uncorrelated variation (dispersion across the major axis) is a measure of cell-to-cell differences in signaling through distinct pathways. We subtract the average intrinsic noise (η^2^(*γ*)) for the gene pairs we are measuring, so we know that the remaining dispersion is due to signaling/pathway noise (η^2^(*P*)) (shown in Fig. 1). Type II experiments also quantify how much cell-to-cell variation is attributable to differences in protein expression capacity (η^2^(G)), which we define as the ability to express and maintain proteins. In the scatter plot of a Type II experiment on the bottom right of Fig. 1, the correlated variation (dispersion along the major axis) is a measure of (η^2^(G)).

### Stoichiometry is Cell Specific, and Signaling Noise is Relatively Constrained in *C. elegans* Intestine Cells

In Type II experiments, we found that cell fate (which ring a cell was in) was the primary determinant of a rigid cell-fate-specific ratiometric setpoint for any given gene pair. Supplementary Fig. S6 shows scatterplots demonstrating how grouping all the cells together artificially inflates signaling noise, compared to grouping cells by ring. So, we grouped cells we measured by the ring from which they came to prevent artificial inflation of (η^2^(*P*)). For the most part, we found that there was little cell-to-cell (and thus, animal-to-animal) relative deviation in signaling through distinct pathways to activate distinct genes.

Fig. 3 shows that there was little signaling noise at the cell level (η^2^(*P*)), relative to intrinsic noise. However, we did notice that some pathways seemed to have more noise; for example there appears to be more signaling noise through *vit-2* than through the *hsp-16.2* pathway. Fig. 3 also shows there is relatively more signaling noise between *hsp-17* and *mtl-2*. This is just not a feature of most reporter pairs we examined. Furthermore, we observed entire populations of animals signal properly. The *hsp-16.2* reporters only expressed after a heat shock and the yolk protein reporters only expressed in intestine cells at the onset of reproductive maturity. For another example of proper signaling, Supplementary Fig. S7 shows an animal resolution experiment demonstrating that, in response to heat shock, animals’ intestine tissue will upregulate eukaryotic translation elongation factor 1 alpha (*eft-3*/*eef-1A*), and downregulate yolk protein production (*vit-2*). Supplementary Figs. S8-S10 show individual scatter plots for Type II experiments and bar graphs of individual cell types. Supplementary Fig. S9 also shows that some cells have relatively more or less noise through some pathways, such as higher signaling noise observed in the intestine cells of ring 1 for *vit-2*, compared to other pathways. Thus, this experimental system is capable of detecting differences in signaling at the cell and population levels, we just did not detect a large degree of signaling noise in young adult intestine cells.

**Figure 3.**
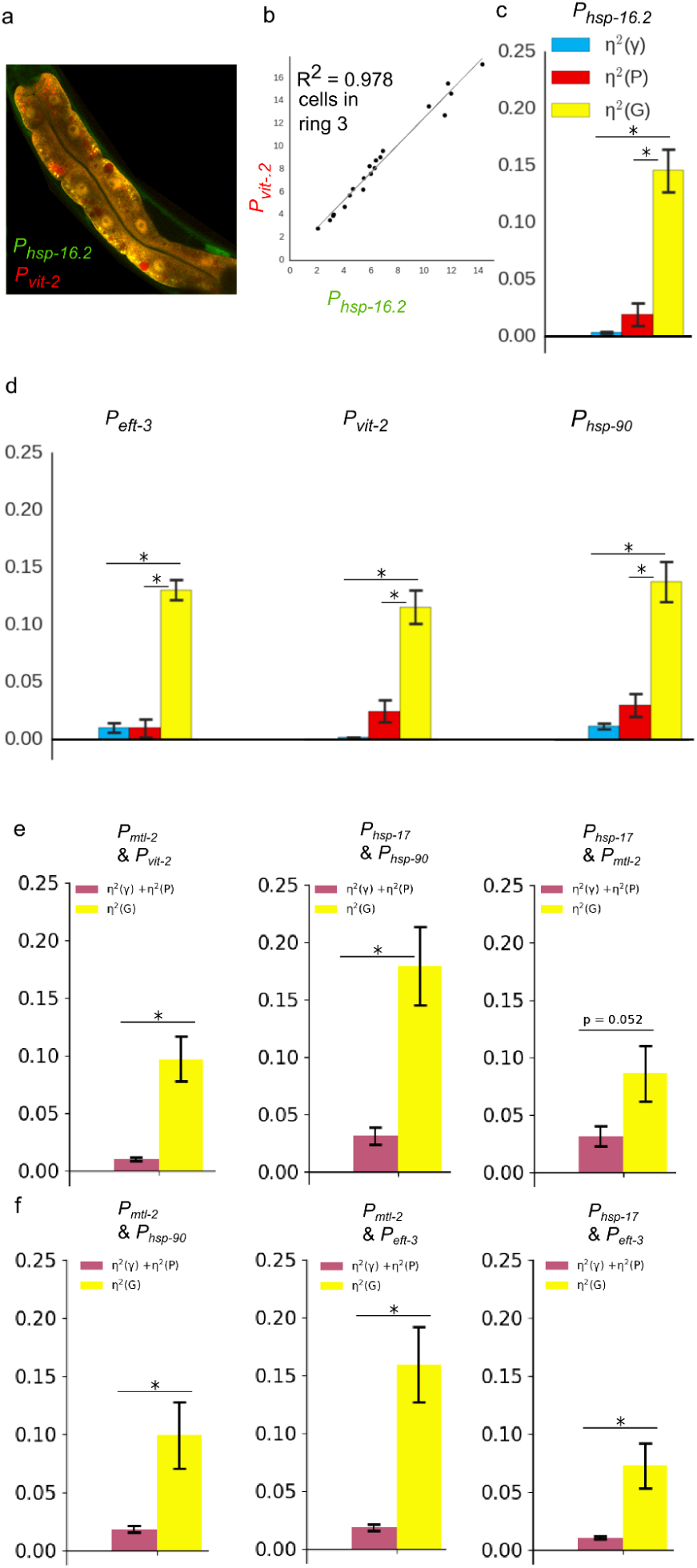
Type II experiments reveal low signaling noise and high variation in protein expression capacity. Variation in protein expression capacity, η^2^(G), is the major source of inter-individual variation in expression. **a.** An example of the animal producing fluorescent reporters (mEGFP and mCherry) from two different promoters (*P*_*hsp-16.2*_ and *P*_*vit-.2*_). **b**. Scatterplot of *P*_*hsp-16.2*_::GFP and *P*_*vit-2*_::mCherry expression in the intestine cells in ring 3. **c**. Average stochastic noise η^2^(γ), variation in pathway activation η^2^(P) and variation in protein expression capacity η^2^(G) for *Phsp-16.2* promoter in cells in the intestine rings 1-4. * indicated statistical significance p<0.05 analyzed by one-way ANOVA with post-hoc Tukey’s HSD test. **d**. Average stochastic noise η^2^(γ), variation in pathway activation η^2^(P) and variation in protein expression capacity η^2^(G) for *P*_*vit-2*_, *P*_*eft-3*_, *P*_*daf-21*_ in the intestine cells in rings 1-4. * indicated statistical significance p<0.05 analyzed by one-way ANOVA with post-hoc Tukey’s test. **e**,**f**. Average correlated and uncorrelated variations for different promoters couples in the intestine cells in rings 1-4. Uncorrelated variation combines stochastic noise of transcription/translation or variable allele access – η^2^(γ) and variation in pathway activation η^2^(P). Correlated variation results from variation in protein expression capacity η^2^(G). * indicates statistical significance *P* < 0.05 by two-tailed t-test; error bars are S.E.M.. For each pair of reporter genes, data are from at least three independent experiments measuring eight intestine cells in at least ten animals per experiment.

### Cell-to-Cell Differences in General Protein Expression Capacity/Protein Dosage Constitute the Largest Intrinsic Axis of Variation in Gene Expression in Intestine Cells

The large (η^2^(G)) components in Fig. 3 shows that we found that most of the variation in gene expression is attributable to cell-to-cell (and thus, animal-to-animal) differences in protein expression capacity (η^2^(G)). Fig. 4e shows (η^2^(G)) visually; two different types of animals expressing two distinct sets of reporter genes from Type II experiments are arranged from dimmest to brightest. Note that the patterns of expression are maintained but that the overall expression level of both reporters is different between animals; to observe this phenomenon, there must be little intrinsic noise (η^2^(*P*)) and little signaling noise (η^2^(*γ*)). We show individual Type II scatterplots and bar graphs organized by cell type and experiment in Supplementary Figs. S8-10.

**Figure 4.**
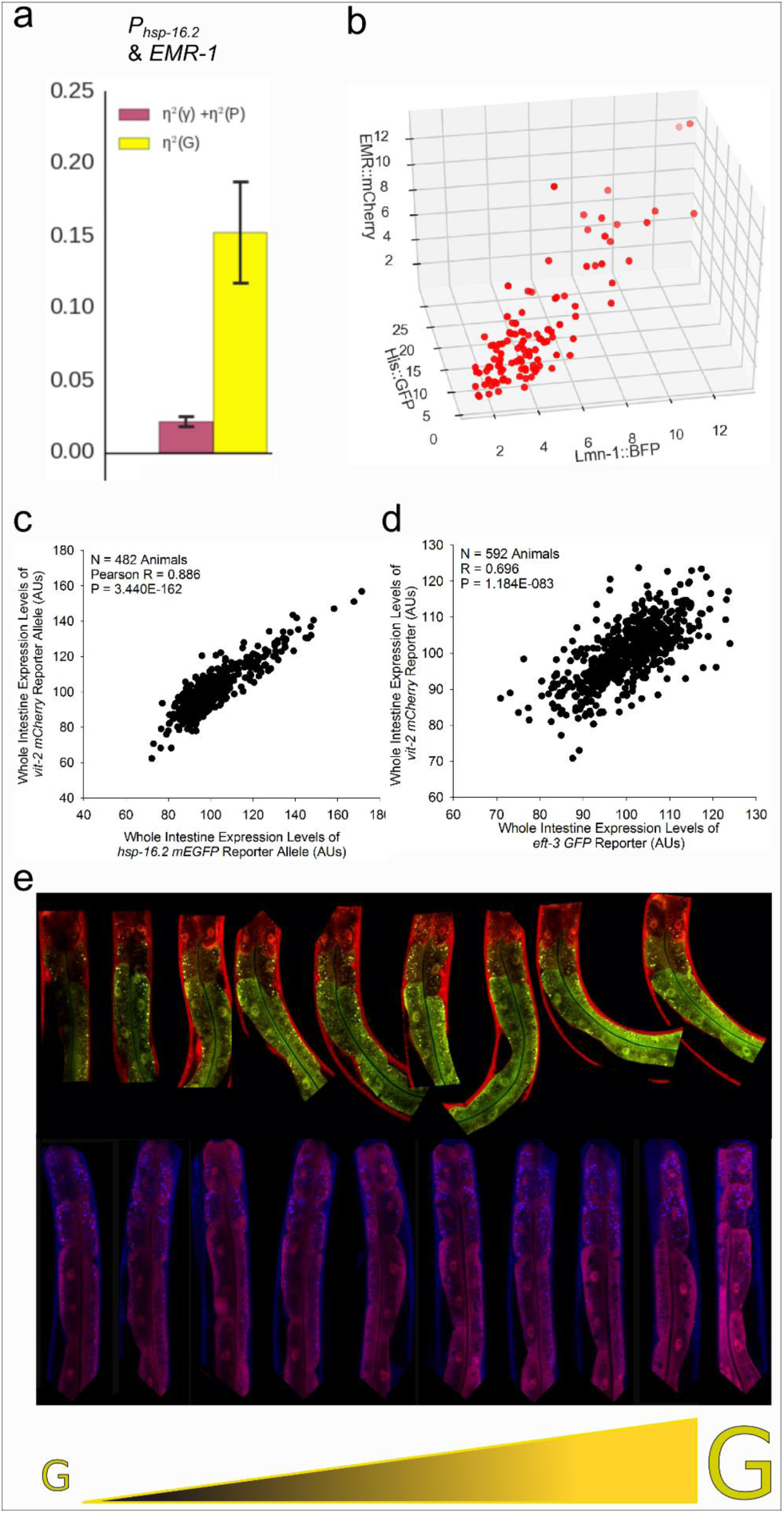
Protein expression capacity is the major axis of variation in gene expression in intestine cells. **a.** Average correlated and uncorrelated variations for *P*_*hsp-16.2*_::mCherry reporter and *P*_*EMR-1*_::EMR-1:GFP fusion protein in the intestine cells in rings 1-4. Uncorrelated variation combines stochastic noise of transcription/translation or variable allele access – η^2^(γ) and variation in pathway activation η^2^(P). Correlated variation results from variation in protein expression capacity η^2^(G). A two-tailed t-test found a significant difference between the two components, *P* < 0.05. **b.** 3-dimensional scatterplot of expression levels of nuclear fusion proteins Lamin:BFP, His2B:GFP and Emerin:mCherry in intestine cells of triple-colored transgenic strain. **c.** Scatterplot of *P*_*hsp-16.2*_::GFP and *P*_*vit-2*_::mCherry expression at whole animal level pooled from three experiments. For pooling, values from each experiment were normalized for mean values. **d.** Scatterplot of *P*_*eft-3*_::GFP and *P*_*vit-2*_::mCherry expression at whole animal level pooled from three experiments. For pooling, values from each experiment were normalized for mean values. **e.** G is a cell nonautonomous, whole tissue property. Multiple pairs of reporter genes covary; animals are generally brighter or dimmer for any pair of reporter genes we measure. Shown are individual animals that vary in expression level for both *P*_*mtl-2*_::*GFP* and *P*_*daf-21*_::*mCherry* (top panel), or, *P*_*eft-3*_::*BFP* and *P*_*vit-2*_::*mCherry* (bottom panel). For the pair or trio of reporter genes measured in cells, data are from at least three independent experiments measuring eight intestine cells in at least ten animals per experiment. For the scatter plots of large numbers of animals we measured at least 100 hundred animals per experiment in three independent experiments to generate the plot shown. For the images, we simply arranged animals from an individual experiment from dimmest to brightest.

### Differences in Protein Expression Capacity Dominate, Even When Measured With Knockins or Fusion Proteins at Distinct Loci in Intestine Cells

These intestine cells express more protein from other, distinctly regulated genes, even if they are on other chromosomes, or fused to other genes (Emerin::GFP; Fig. 3e). We wanted to confirm our findings with additional fusion proteins. We also wanted to measure fusion proteins that would not have signal convoluted by abstract spatial patterns and autofluorescence. Therefore, we chose to quantify proteins that localized to the relatively low autofluorescence, and entirely segmentable, nucleus. We quantified a His2B knockin in conjunction with Emerin and Lamin fusion proteins in the nuclei of animals’ intestine cells. We found that there was significant covariation of these proteins among the cells of animals, with the major axis of variation being the general abundance, shown in a 3D scatter plot in Fig. 4; additional experiments shown in Supplementary Fig. S11.

### The Intestine Tissue Varies Most in General Protein Expression Capacity, Even in Large Populations of Animals

We examined large populations of animals to confirm differences in general protein dosage constitute the major axis of animal to animal variation, as we saw at the cell level. When we examined distinctly regulated reporters in hundreds of animals, we found that animal-to-animal variation in G was dominant. When we tried to quantify covariation between signals in unconstrained animals in the COPAS Biosort, or in a microfluidic, image-based quantification device we developed^24^, we found little covariation between genes we know to be highly correlated inside individual intestine cells. This appears to be due to the variable positioning of the animals in front of the detectors causing an uncoupling of signals. These higher throughput devices will get the right answer for the average expression level of the two different reporters, but will not get the right answer for covariation. Animals need to be tightly restricted. Indeed, as we show in Fig. 4, the same trend for differences in G can be seen, even at animal resolution, provided animals are constrained and a tissue specific marker is used to extract relatively purer signal from a focused region of interest (the intestine tissue). Signals from other tissues differentially contributed to the intestine signal we were trying to measure using an epifluorescent scope to examine larger populations of animals. We believe the deviations from the trendline with the *eft-3* and *vit-2* reporters are from out of plane non-intestinal signals contributing to the intestine signal. We think the *vit-2* and *hsp-16.2* reporters are more representative of what we see at the cell level because both reporters are mostly expressed in the intestine (*vit-2* exclusively so), whereas *eft-3* signal is also quite strong in muscle and hypodermal tissues surrounding the intestine.

### Increased Protein Expression Capacity is Underlain by Both Increased Production and Maintenance of Protein in Normal and Heat Shocked Conditions

We wanted to test the hypotheses that high gene expression capacity was due to either better protein production or better maintenance/decreased turnover. Therefore, we tested to see if animals that made more fluorescent protein did so by increased production or decreased turnover in both normal and heat shocked conditions. We quantifying the age of protein *in vivo* using a fluorescent timer protein that matures from green to red in about 48 hours^25^. We controlled the reporter with the *eft-3* promoter, which is both constitutive and most highly correlated with both *hsp-16.2* and *hsp-90* chaperone reporters. We found that animals that are brighter or dimmer have similar ratios of both new and old protein (e.g., exactly 2.71 new to old ratio in both the top and bottom 10% of heat shocked animals). Supplementary Fig. S12 shows the trend line is linear and the animals at the top and bottom do not deviate. Consistent with the dual roles of chaperones in protein production and maintenance, we find that animals that are brighter are better at both protein production and maintenance to a greater degree than dimmer animals. Supplementary Fig. S8 also shows that this timer protein does indeed detect a higher proportion of relatively older protein in relatively older animals.

### Additional Corroborating Data

As previously reported^10,11,26^, we also found that chaperone or chaperone-related reporters can correlate with or predict biological outcomes. Prior work showed that chaperones increase the activities of loss of function mutations (but not nulls!)^10^, suggesting that gain of function mutations would also sometimes have higher activities with higher abundance of some chaperones. Thus, we extended previous work with loss of function mutations to include alleles that conferred gains of function. In Supplementary Material Section 3: Additional Correlations Between Phenotypes and Reporter Genes, we show that adult expression of the *hsp-16.2* biomarker correlates with the penetrance of a Ras gain of function mutation that acted during larval development in Supplementary Figs. S13&S14. We also show that the expression of the highly chaperone-correlated *eft-3* reporter in L1 larvae predicts growth to adulthood on neomycin in animals also expressing a NeoR gene (gaining the function of neomycin resistance) in Supplementary Fig. S15. In Supplementary Material Section 4: Persistence of States, we examine evidence of persistence, finding that the embryonic state of high yolk load or the L1 larval state of high gene expression (discerned via *eft-3* reporter levels) are not correlated with adult gene expression levels, shown in Supplementary Fig. S16. In Supplementary Fig. S17 we show a type of larval variation in gene expression that we do not see in adults that will require further study to understand how it fits in the context of developmental progression and other kinds of noise in gene expression.

### Working models for how intestines may end up in high protein dosage states after heat shock

The natural variation in the *C. elegans* chaperone system is a property that is consequential, and likely selected for over geological time, if we subscribe to the prior conclusions of Klaus Gartner when he was pondering the origins of nongenetic variation in murine systems^27^; it seems reasonable to do so. In our working model, animals with higher concentrations of the *hsp-16.2* lifespan/penetrance biomarker in their intestine cells have higher concentrations of other proteins in those cells. These differences can arise from epigenetic differences^28^ and seemingly stochastic environmental perception differences^14,15^.

Given the known role of chaperones in protein production, maintenance and turnover, we speculate that chaperone levels may sometimes determine differences in global protein dosage^6^. Figs. 5a&b show a working model of how variation in adult intestinal protein expression capacity may arise. In this model, differences in environmental perception result in differences in AFD sensory neuron firing^14,29,30^, which results in differences in insulin signaling in the peripheral intestine tissue^14,29,30^, causing an upregulation of chaperones^31^. Fig. 5c shows a general model of how hypomorphic and hypermorphic protein activities may be increased or decreased by chaperones (via protein expression capacity) to cause differences in the manifestation of traits.

**Figure 5.**
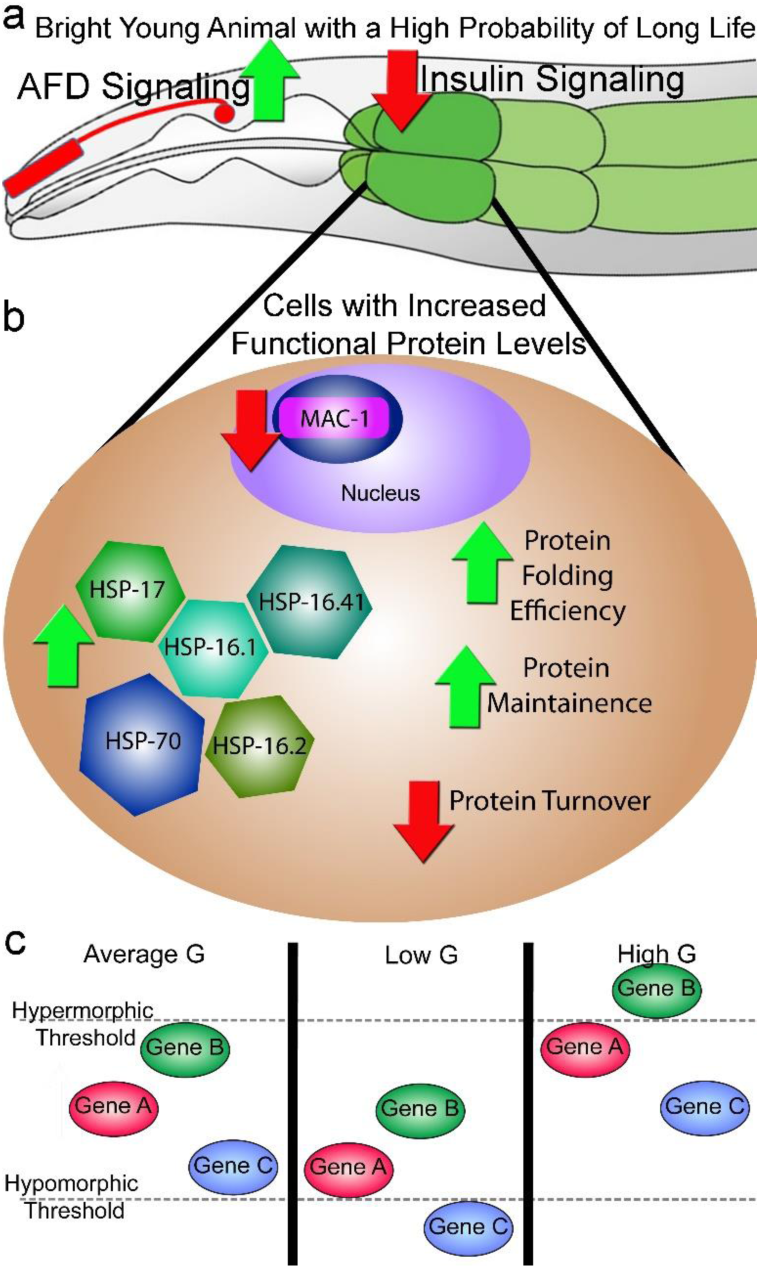
Working Model. **a.** Genetic experiments from Mendenhall et. al. show that interindividual variation in the *hsp-16.2* reporter gene that is a biomarker for mutation penetrance and lifespan arises from differences in insulin signaling and depolarization of the AFD neuron pair. Thus, animals with low insulin signaling and high neuronal depolarization would have the highest expression of the *hsp-16.2* lifespan/penetrance biomarker and have the longest lifespans, highest penetrance of hypermorphic phenotypes and lowest penetrance of hypomorphic phenotypes. **b.** Diagram of the molecular and physiological differences in cells that underlie global differences in protein dosage. Solid arrows are directly supported by experiments in *C. elegans* using reporter genes (here in Figs. 3,4 & S12 and previously^28^) and microarrays (previously^28^). **c.** Three panels showing consequences of different states of protein expression capacity (G). Left panel shows an individual with average G and wild-type phenotypes. Middle panel shows an individual with low G and penetrance of a hypormorphic phenotype for the allele of Gene C. Right panel shows an individual with high G and penetrance of a hypermorphic phenotype for the allele of Gene B.

In distinct interpretations of the working model, there may, or may not, be trade-offs between somatic fitness and reproductive fitness, and, these trade-offs can manifest distinctly in terms of how they affect protein production or export from the intestine. In the first model interpretation, high chaperone/*hsp-16.2* biomarker expressing animals respond better to heat shock by making and maintaining protein better, they live longer, and there is no trade-off with progeny production, there are just winners and losers. This is most supported by the data. In another model interpretation, there is a trade-off, both bright and dim animals produce and maintain similar amounts of protein, but the dimmer animals simply export more protein to their oocytes, effectively atrophying the intestine faster as in^32^. In another version of the trade-off model, the dimmer animals do make more protein to make more progeny, consistent with a trade-off after heat shock^10^, and the brighter animals have more protein because of increased maintenance, and not better production. It is interesting to notice in this regard, that we have previously found that animals expressing high levels of *hsp-16.2* biomarker have lower levels of *mac-1*/*rix7p* ribosome export factor. It is therefore intriguing to speculate that bright animals synthesize less protein and simply maintain mature proteins better, as that could be due to high levels of chaperons and low insulin signaling. Accordingly, decreased insulin signaling is associated with decreased global protein turnover^33-35^. Yet, this is also not supported by the current data, as animals that made more timer protein under *eft-3* promoter control had the same ratio of relatively younger versus relatively older protein (Fig. S12). It will be interesting in the future to determine these molecular mechanisms underlying the states of high and low global protein dosage. These alternative interpretations are plausible, but not currently supported by the data. Supplementary Material Section 5: Trade-offs and Supplementary Fig. S18 show that we can detect trade-offs in other biological scenarios, we just did not after adult heat shock^28^.

## DISCUSSION

We examined sources of variation in gene expression in intestine cells in *C. elegans*. Our results show that intrinsic noise and signaling noise are minimal, and that the major contribution to differences in gene expression is differences in general protein expression capacity/effective protein dosage. This axis of variation is dominant, whether animals are heat shocked or not. It is dominant in large or small populations of animals. Observations from reporter proteins, fusion proteins and phenotype are all consistent with the idea that expression of significant portions of the proteome covaries with expression of chaperones. We found that differences in protein expression capacity can be consequential in early larval development, but do not appear to persist into adulthood (at least not from the L1 diapause), consistent with a prior report showing embryonic physiology (yolk protein amount) did not correlate with adult physiology (lifespan)^36^. We found that high G animals maintain significantly higher fractions of both old and young protein than low G animals, with or without heat shock. Thus, the animals responding better to heat shock are better able to produce and maintain protein in their intestines. Given the recent finding that the intestine consumes itself ^32^, the 2011 study wherein Sanchez-Blanco et al showed that most reporter genes they measured in the intestine predicted lifespan seems more logical^21^; more remaining protein to burn equated to more lifespan.

### Chaperons and mechanisms of variation in proteome dosage

Given the role that chaperones play in protein production, maintenance and turnover, it is not surprising that chaperone reporters predict differences in biological outcomes, from penetrance of mutations to lifespan^6^. What was initially surprising was the covariation of expression of these reporters with other distinctly regulated genes. This fact can be explained if expression of these genes is affected by differences in gene expression capacity. Though, again, in retrospect, given the role of chaperones, it should not have been surprising. And, given that metazoans have to coordinate the activities of multiple cell types in multiple tissue types, it may not be surprising that signaling noise was incredibly restricted, relative to the one other single celled eukaryote in which these kinds of measurements have been made.

Another source of variation in protein dosage could be variation in genome dosage. Indeed, our previous work showed that animals that were heterozygous for the biomarker expressed about half as much reporter protein – so, like other systems, gene dosage affects expression level in *C. elegans*. However, due to technical limitations of our fixation procedure, we were not able to reliably make simultaneous measurements of fluorescent protein and nuclear genome content to determine any correlation between ploidy and gene expression level. We did not pursue this further because, currently, we do not think variation in intestinal ploidy drives variation in intestinal proteome dosage for two reasons. First, we see that the variation in biomarker expression is controlled by AFD neuron depolarization^14^, which has no known role in ploidy control, nor does the *gcy-8* mutant we used to shut down AFD depolarization have the ploidy-related smaller body phenotype ^37^. Second, in other work focused on cis control of intrinsic noise, we found that diploid muscle cells have about the same amount of extrinsic noise (revealed here as G) as polyploid intestine cells^38^.

The high correlation of chaperone reporters with phenotypes (Figs. S13-15 and ^10,12,13,26^), distinctly regulated transcriptional reporters (Fig. 3), fusion proteins (Fig. 4) and knockins (Fig. 4) suggests that at least some significant fraction of cellular proteome covaries in some circumstances. This is worth considering in the larger context of any biological scenario. However, we do not yet know what fraction of the cellular proteome for which protein dosage correlates with chaperone abundance. While our experimental system did detect gene expression levels changing in response to external signals (Fig. S7), and instances of visually detectable intrinsic noise and signaling noise, for the most part, the small fraction of the genome for which we examined expression levels or phenotypes covaried fairly well (e.g., Figs. 3&4). Presumably, additional work exploring variation in the expression or activity of the myriad of remaining genes will identify additional axes of variation that will help define specific fractions/modules of the proteome that covary in different scenarios. For example, Sanchez-Blanco et al found another, uncorrelated axes of variation using a fluorescent protein reporters^21^. While most reporters they looked at in eight day old adult worm intestines were highly correlated (e.g, r = 0.4-0.87), one single reporter, controlled by the promoter for *C26B9.5*, which encodes a serine protease, did not correlate well^21^. Thus, while we did not see additional major axes of uncorrelated variation, it may be possible to find other axes of G, or significant differences in P, with additional fluorescent reporter genes or fluorescent knockins.

### Mechanisms of cell-to-cell variation in gene expression in other tissues

Other reports have shown that intrinsic noise can be a significant component of cell-to-cell variation in gene expression. In yeast, intrinsic noise was an order of magnitude higher than in worm intestine cells^19^. In human cell culture, intrinsic noise, to the point of monoallelic gene expression, has been reported quite frequently now^39^. We believe that intrinsic noise of gene expression will be a larger component of cell-to-cell variation in gene expression in diploid tissues. Indeed, in other work, we found that intrinsic noise is significantly higher in diploid muscle cells, compared to the polyploid intestine cells examined here^38^. Furthermore, understanding this can kind of noise in gene expression may have implications for understanding how people can live with and escape the consequences of dominant negative mutations, like some oncogenes^40^. It is worth noting that the phenotypes predicted by the chaperone reporters have to do with the action of genes in the polyploid intestine and hypodermis^10,11^. This is true for the Ras gain of function affecting vulva development reported here and at least partially true for the more organismic phenotypes of thermotolerance, lifespan and neomycin resistance, which must rely on the intestine and hypodermis, at least in part.

### Chaperone variation in health and disease

Natural or pharmaceutical manipulation of chaperone systems in different tissues has the potential to affect both health, in terms of robust living, and disease, in terms of affecting the activity of specific disease alleles. Anti-HSP90 drugs have been used to attempt to attenuate the activity of Ras gain of function mutations in cancer treatments for almost a decade^41,42^. Unfortunately there are side effects when targeting the master regulator of the heat shock response; for example, the HSP90 inhibitor AUY922 had the side effects of diarrhea, skin rash, hyperglycemia, and night blindness^43,44^. It may be that other components of the protein chaperone system can be targeted with less collateral damage than targeting the master regulator of the heat shock response. Alternatively, increasing the activity of some genes or the dosage of large portions of the proteome may be desirable in other biological scenarios. For example, increasing the dosage of chaperones through increasing the abundance of individual chaperones^45^ or the master regulatory transcription factor, HSF-1^46,47^, has been shown to increase lifespan in *C. elegans*. Thus, designing small molecule therapies that elicit specific chaperone responses affecting specific subsets of traits may be a worthwhile endeavor for improving human health.

### Differences in protein abundance/activity and missing heritability

Natural variation in chaperone subsystems affects the penetrance and expressivity of some traits, and is therefore likely to be responsible for the missing heritability of some traits. We know distinct chaperones affect the manifestation of distinct sets of traits^10^. We know that there is an epigenetic component to heritability^48^. And we know that chaperone levels can be epigenetically heritable^28^. We now know, at the resolution of single cells, operating at the protein level, for one metazoan tissue, that differences in the dosages of many proteins correlate with and are likely influenced by natural variation in chaperones. So, it may be that the current inability of genetic variants to account for differences in the manifestation of traits is due to another factor accounting for such differences, such as intrinsic, heritable physiological variation in chaperone levels.

## Acknowledgements

Some strains were provided by the CGC, which is funded by NIH Office of Research Infrastructure Programs (P40 OD010440). We thank the other laboratories and scientists that have contributed to the understanding of cell-to-cell variation in gene expression not referenced in this manuscript; a complete literature review on non-genetic, non-environmental biological variation was beyond the scope of this study. We thank George Martin for thoughtful discussions about the manuscript. We thank the Lehner lab for providing their *vit-2::GFP* knockin. We thank the Goldstein Lab for providing their *his-72::GFP* knockin.

## Author Contributions

AM, NB, TEJ and RB designed the study. NB, AM and RB analyzed and interpreted the results. AM, RB, TEJ and MK provided funding. BS, NB, PT, AM and SY generated DNA and transgenic nematode strains. AM and NB conducted the imaging experiments and performed image cytometry. AM, PT and BS conducted the flow sorting and phenotyping experiments. NB and AM analyzed the raw data. AM, RB and NB wrote the initial manuscript. RB, MK, TEJ, PT, BS, SY, AM and NB reviewed and revised the manuscript.

## Data Availability Statement

The datasets generated during and/or analysed during the current study are available from the corresponding author on reasonable request.

