## Supplementary Information for "*hsp-16.2* chaperone biomarkers track physiological states of proteome dosage"

**for**

**by**

**Burnaevskiy et al.**

### Table of Contents

Figures S1-S12........................................................................................Pages 2-15

Supplementary Material Section 1: Materials and Methods.........................Pages 16-21

Supplementary Material Section 2: Analytical Framework…………………..Pages 22-31

Supplementary Material Section 3: Additional Correlations between Phenotypes and Reporter Genes………………………………………………………………..Pages 32-38

Supplementary Material Section 4: Persistence of Physiological States…..Pages 39-42

Supplementary Material Section 5: Trade-offs………………………………..Pages 43-45

References………………………………………………………………………..Page 46

**Supplementary Figure S1. Cartoons Depicting Intrinsic Noise States.** A Cartoon Diagram showing how we generate animals for Type I experiments to examine intrinsic noise is shown. Top section shows two homozygous animals being bred together to create an archetypal hybrid. The archetypical hybrid has a stereotyped, anterior-posterior pattern of expression, which we do observe for all combinations of reporters in intestine cells. Bottom Panels show (left) what low differences in γ would look like, (center) what cell nonautonomous differences in γ would look like, and (right) what cell nonautonomous differences in γ would look like.

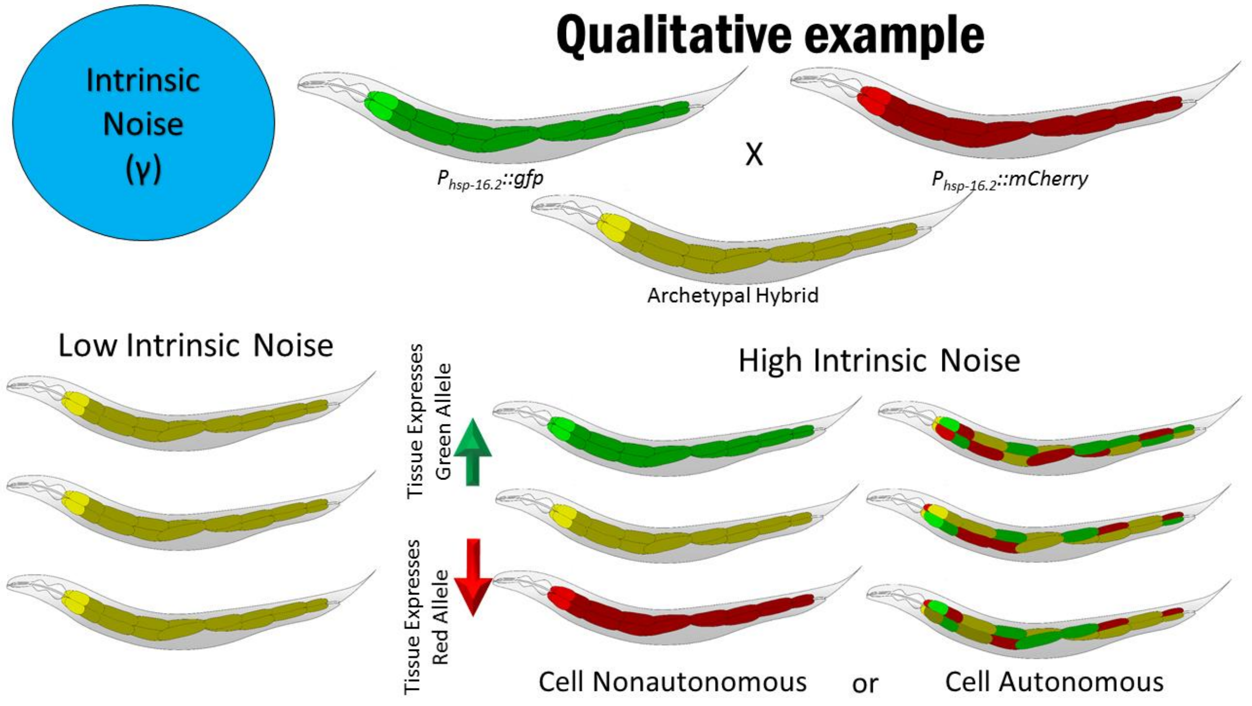

**<u>Supplementary Figure S2. Cartoons Depicting Signaling Noise States.</u>** A Cartoon Diagram showing how we generate animals for Type II experiments to examine signaling noise is shown. Top section shows two homozygous animals being bred together to create an archetypal hybrid. The archetypical hybrid has a stereotyped, anterior-posterior pattern of expression, which we do observe for all combinations of reporters in intestine cells. Bottom Panels show (left) what low differences in P would look like, (center) what cell nonautonomous differences in P would look like, and (right) what cell nonautonomous differences in P would look like. All of these images are assuming intrinsic noise is constrained. Cell autonomous signaling noise would experimentally look like intrinsic noise, which is why it is critical to establish some measures of intrinsic noise before attempting to precisely quantify signaling noise.

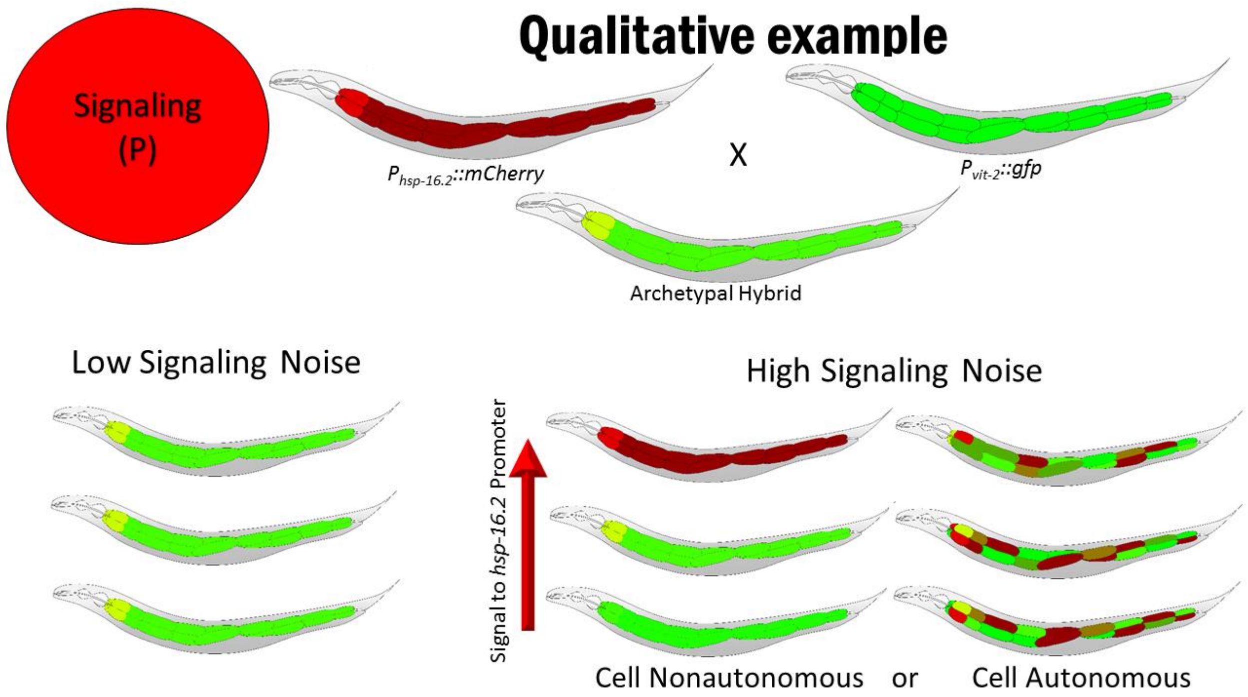

**Supplementary Figure S3. Cartoons Depicting Protein Expression Capacity States.** A Cartoon Diagram showing how we generate animals for Type II experiments to examine gene expression capacity is shown. Top section shows two homozygous animals being bred together to create an archetypal hybrid. The archetypical hybrid has a stereotyped, anterior-posterior pattern of expression, which we do observe for all combinations of reporters in intestine cells. Bottom Panels show (left) what low differences in G would look like, (center) what cell nonautonomous differences would look like (a cartoon approximation; see Fig. 4 for actual images), and what cell nonautonomous differences in G would look like. All of these images are assuming intrinsic noise and signaling noise are constrained. The cartoons are intended to show the correct ratiometric setpoints, but different absolute values for the whole intestine for cell nonautonomous and for individual cells that randomly vary in G, but keep the right ratio of expression in the cell autonomous G cartoons.

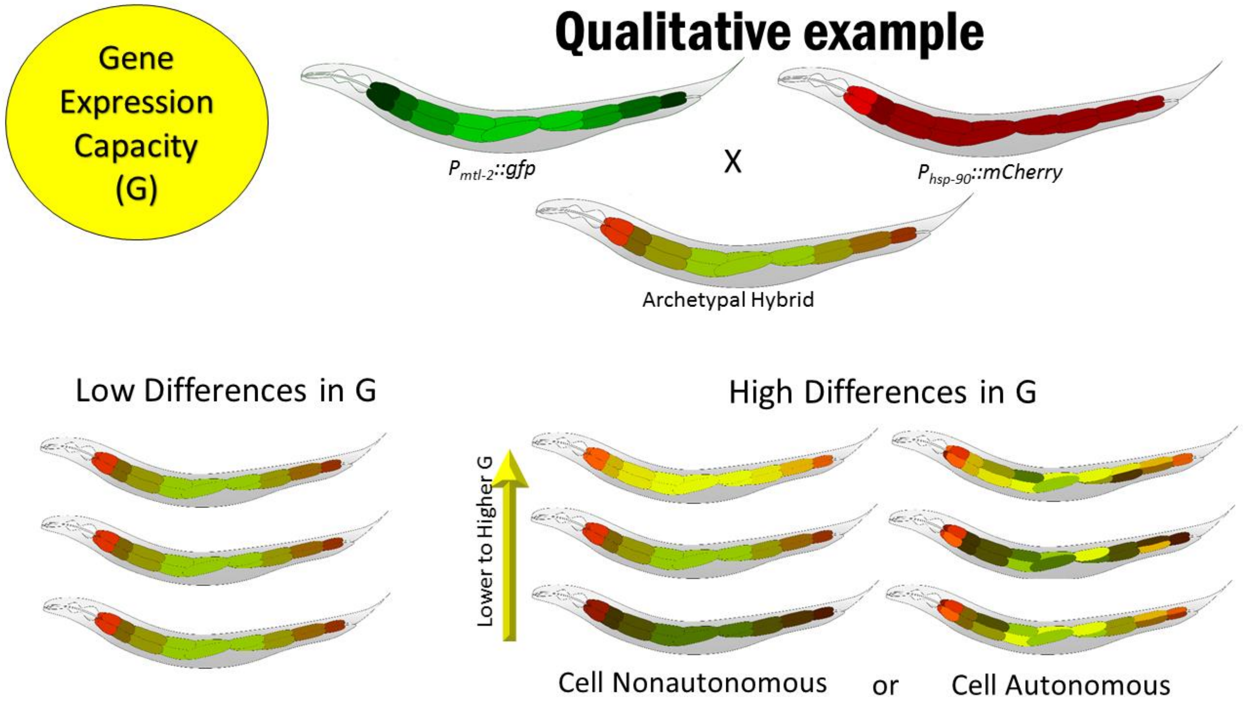

**Supplementary Figure S4. Type I experiments Supplementary scatterplots.**

Scatterplots of two identical promoters with different fluorescent protein outputs expressed from homologous chromosomes (type I experiment). Far left scatterplots show all cells measured in a given experiment. Scatterplots on the right show expression of reporters in the cells from particular intestine rings. One of at least three repetitions per reporter gene pair is shown.

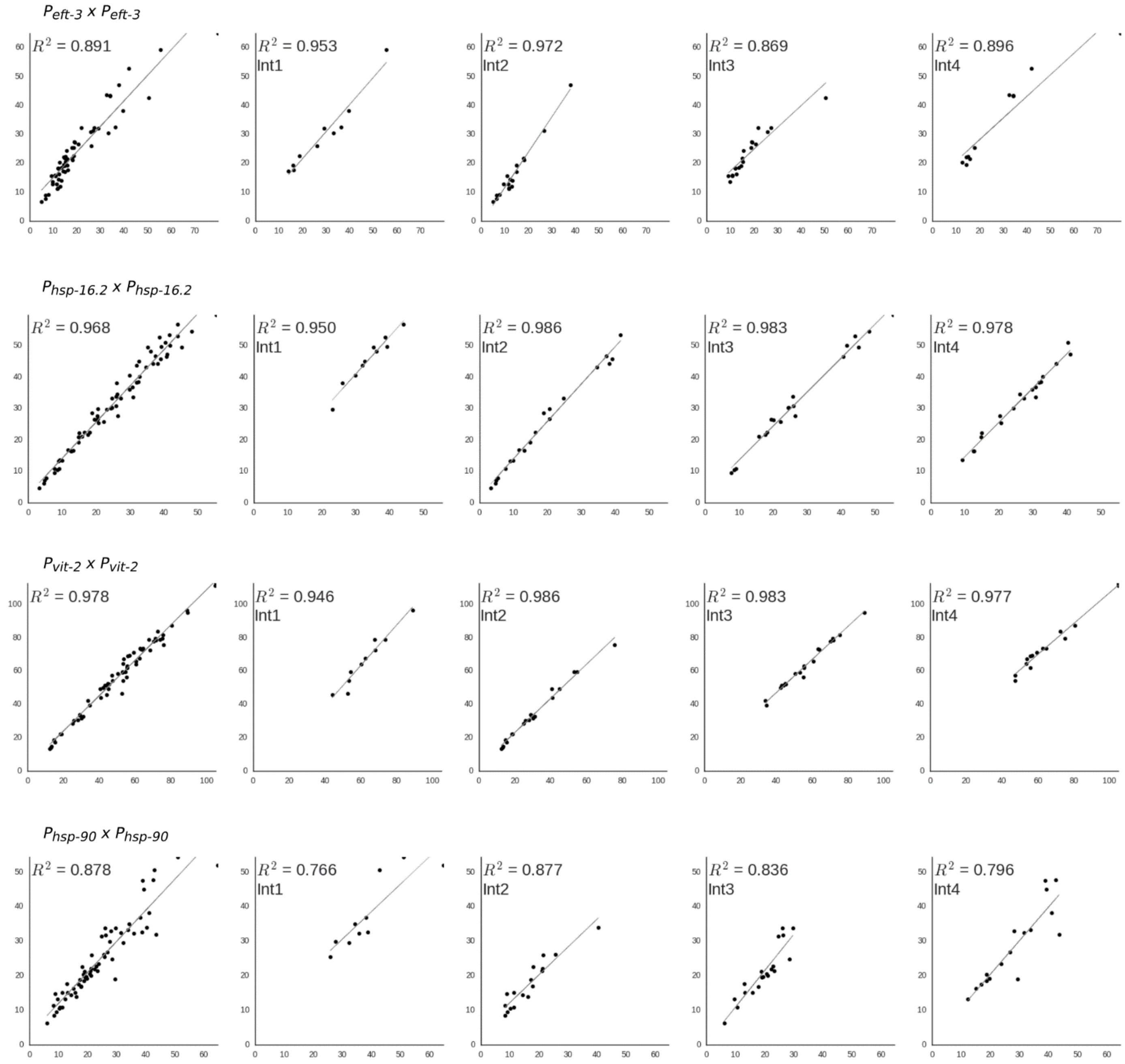

**Supplementary Figure S5. Type I experiments Supplementary bargraphs.**

Correlated and uncorrelated variation for $P_{hsp\text{-}16.2}$, $P_{eft\text{-}3}$, $P_{vit\text{-}2}$ and $P_{\text{-}hsp\text{-}90}$ based reporter genes in intestine cells in rings 1-4. In type I experiment uncorrelated variation arises from stochastic noise of transcription/translation or variable allele access – $\eta^2(\gamma)$; correlated variation is a combined result of variation in gene expression capacity $\eta^2(G)$ and variation in pathway activation $\eta^2(P)$. This bar graph consists of data for each cell type from at least three independent experiments quantifying two cells per ring in each of at least ten animals (at least 60 data points).

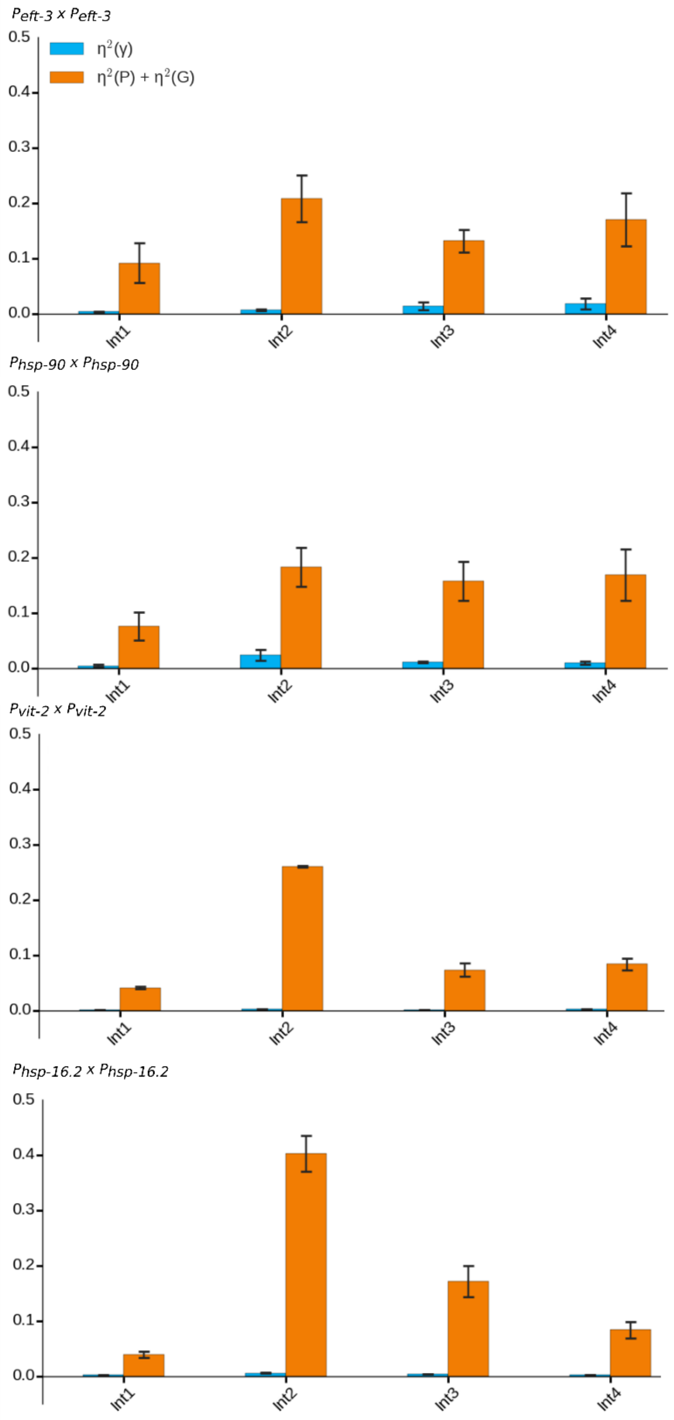

**Supplementary Figure S6. Evidence for splitting cells by fate for analysis in type II experiments.**

Scatterplots of expression of two reporters from the different reporter genes, $P_{hsp\text{-}17}$ and $P_{mtl\text{-}2}$, grouped by ring (one through four) or combined in one plot (right panel). Cell fate determines ratiometric setpoint for expression of two distinct genes. When cells are not split by fate, correlation of expression is quite low. Correlation of expression is much higher when cells are grouped by fate. Data shown is from an independent experiment quantifying the aforementioned reporters in intestine cells located in intestine rings one through four.

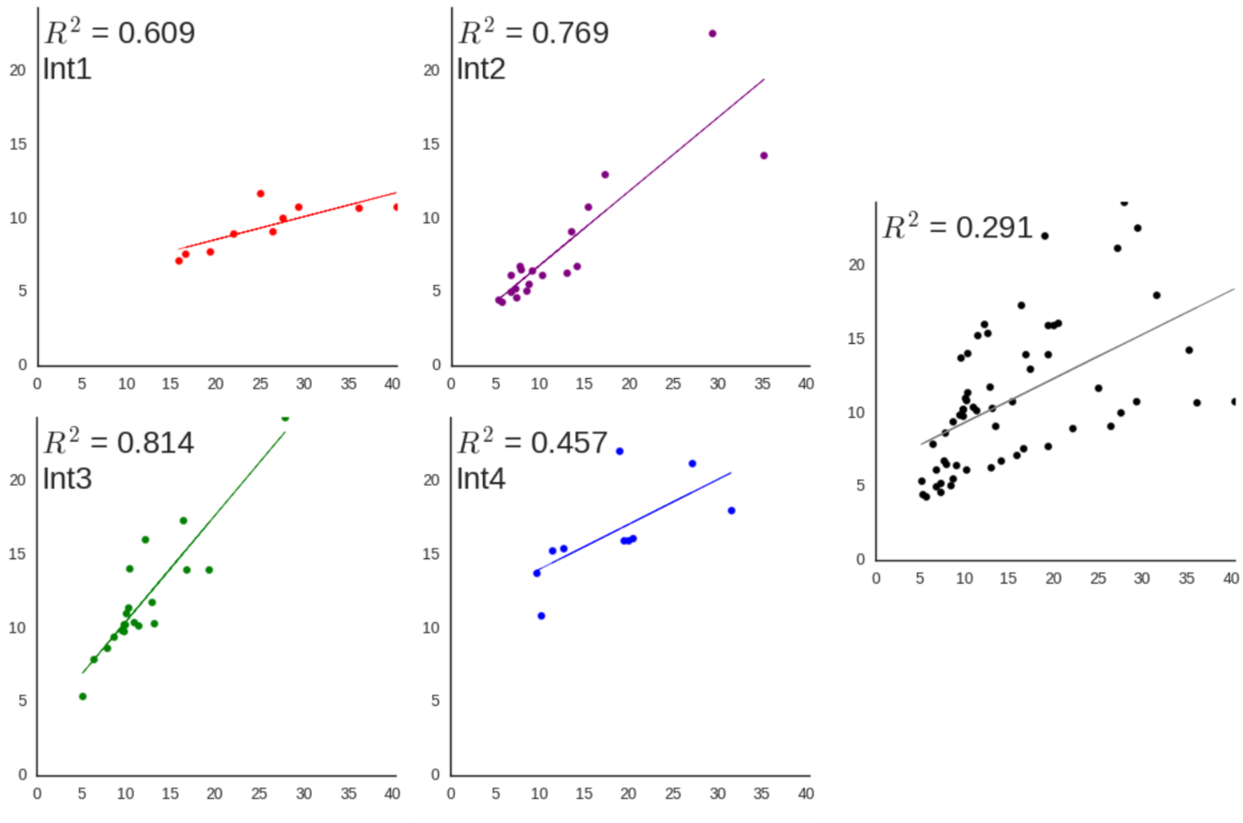

**Supplementary Figure S7. Evidence for proper signaling.** Scatterplot of $P_{eft\text{-}3}$::GFP and $P_{vit\text{-}2}$::mCherry expression at whole animal level with and without heat shock. For heat shock, day 1 adults were incubated for 1hr at 35°C and imaged 24 hours later together with age-matched non-heatshocked animals. a. A scatter plot of signaling changes in *vit-2* and *eft-3* reporter genes in response to heat shock from an individual experiment is shown. b. Bar graphs quantifying average expression levels from individual experiments are shown; error bars are standard error of the mean. Animals were measured with an epifluorescent microscope at animal resolution, mounted on cover slips as described in methods. At least 30 animals were used per group per experiment.

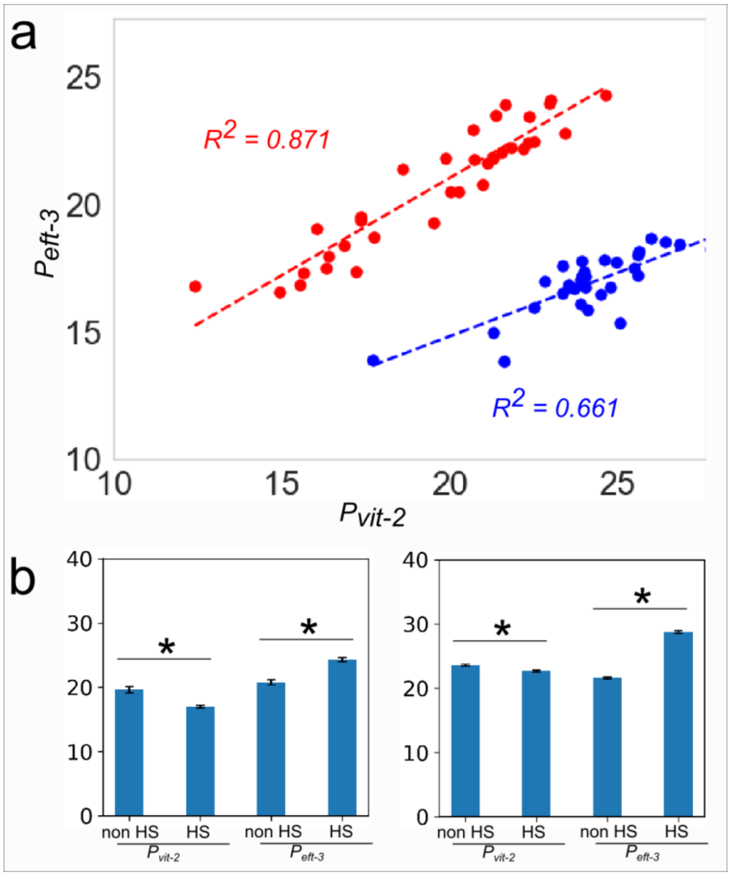

**Supplementary Figure S8. Type II experiments Supplementary scatterplots.**

Scatterplots of expression of two distinct reporter genes (type II experiment). Far left scatterplots show all cells measured in a given experiment. Scatterplots on the right show expression of reporters in the cells from particular intestine rings. One of at least three repetitions per reporter gene pair is shown.

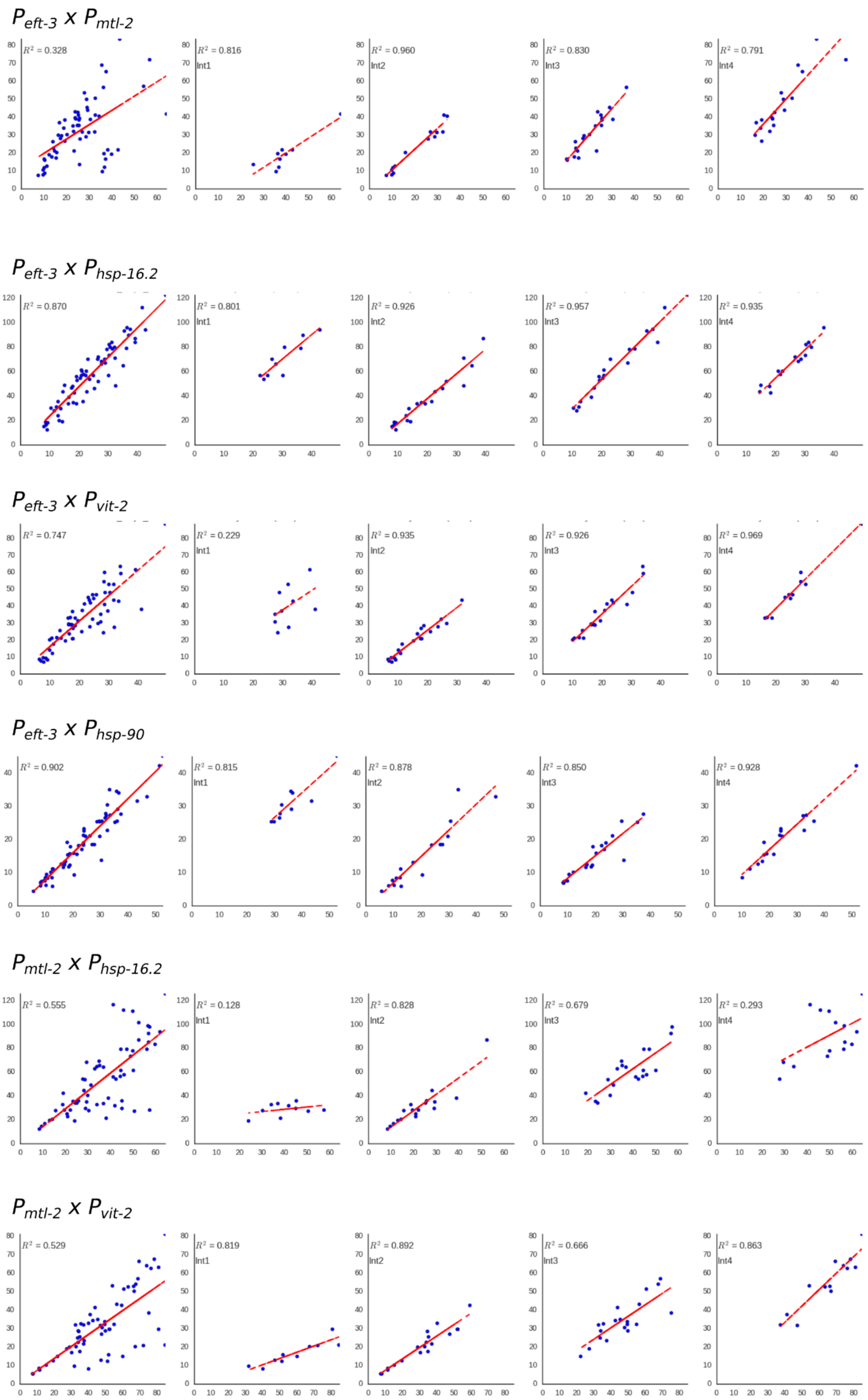

Type II experiment scatterplots Part II

$P_{mtl\text{-}2}$ x $P_{hsp\text{-}90}$

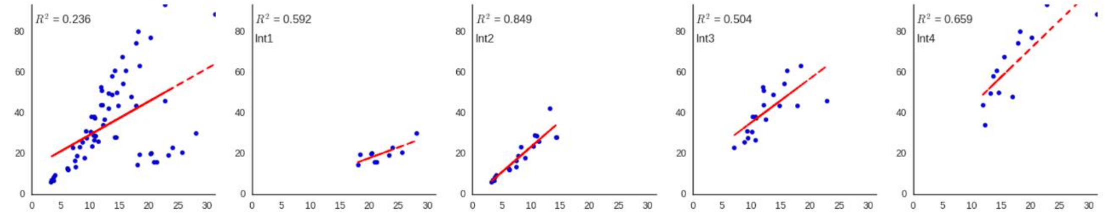


$P_{hsp\text{-}16.2}$ x $P_{vit\text{-}2}$

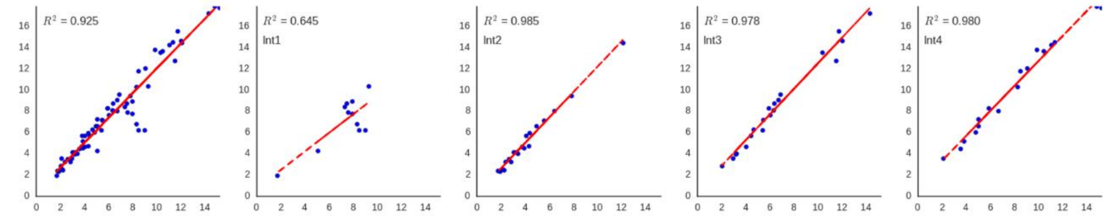


$P_{hsp\text{-}16.2}$ x $P_{hsp\text{-}90}$

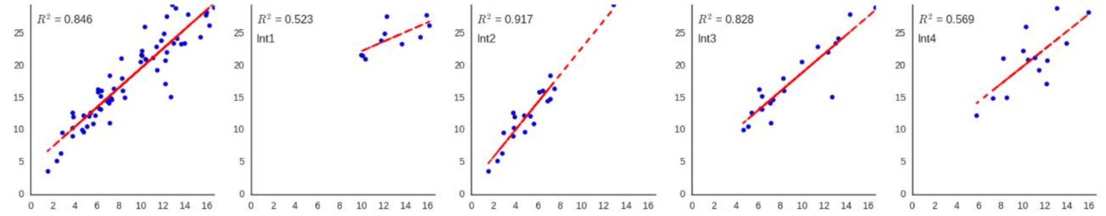


$P_{hsp\text{-}90}$ x $P_{vit\text{-}2}$

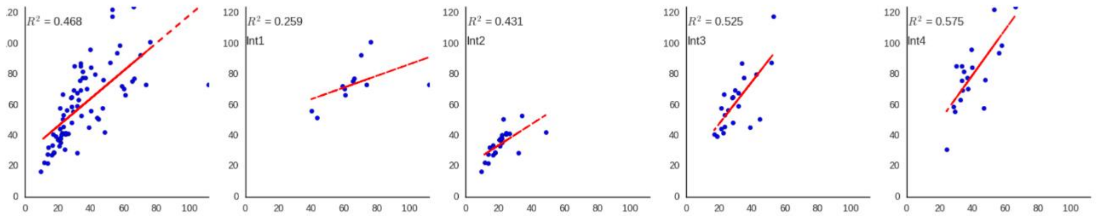


$P_{hsp\text{-}17}$ x $P_{mtl\text{-}2}$

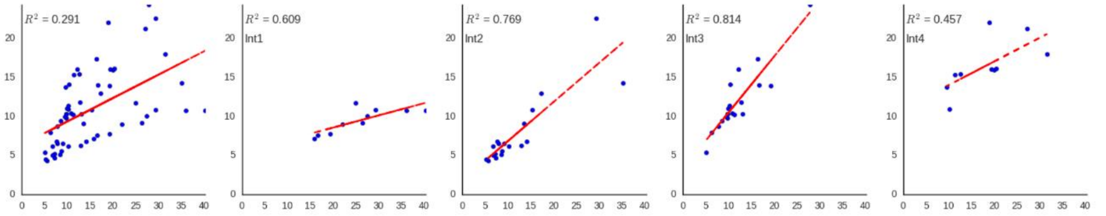


$P_{hsp\text{-}17}$ x $P_{hsp\text{-}16.2}$

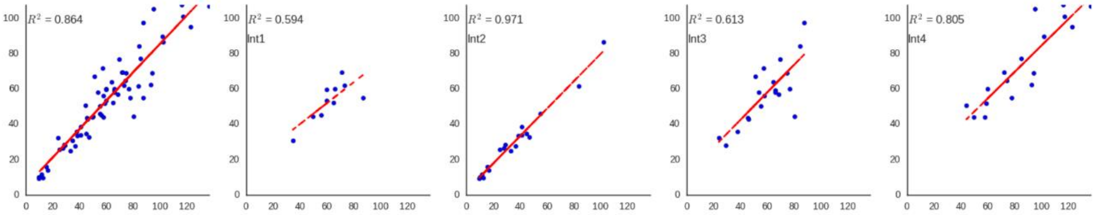

Type II experiment scatterplots Part III

$P_{hsp\text{-}17}$ x $P_{vit\text{-}2}$

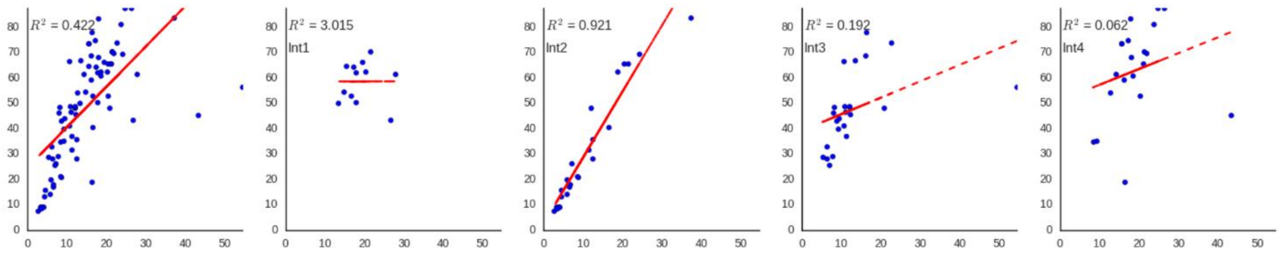


$P_{hsp\text{-}17}$ x $P_{hsp\text{-}90}$

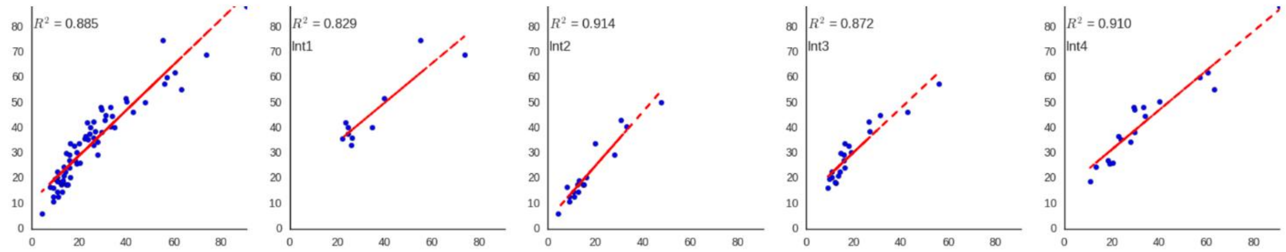


$P_{hsp\text{-}17}$ x $P_{eft\text{-}3}$

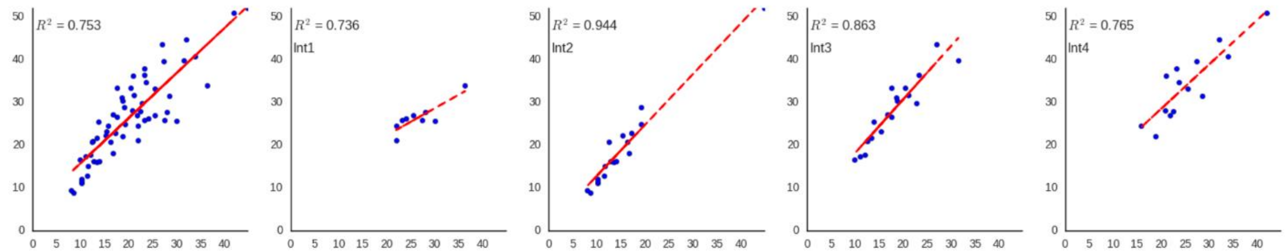


EMR-1 x $P_{hsp\text{-}16.2}$

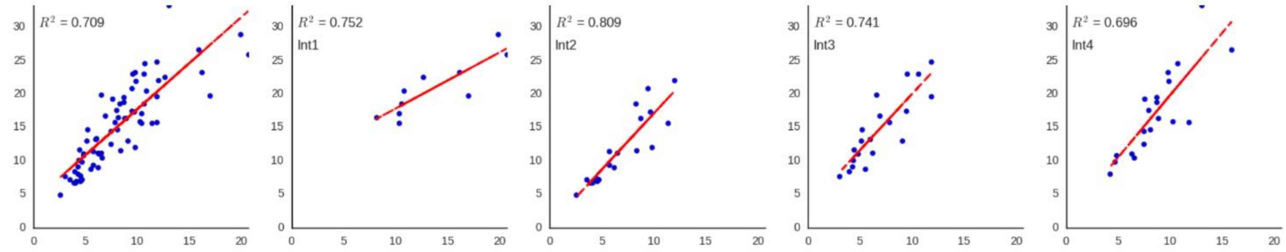

**Supplementary Figure S9. Type II experiments Supplementary bargraphs.**

Stochastic noise $\eta^2(\gamma)$, variation in pathway activation $\eta^2(P)$ and variation in gene expression capacity $\eta^2(G)$ for *P<sub>hsp-16.2</sub>*, *P<sub>vit-2</sub>*, *P<sub>eft-3</sub>* and *P<sub>hsp-90</sub>* for cells in intestine rings 1-4. These bar graphs are composed of data from three independent experiments quantifying expression from two cells per ring per animal for at least ten animals per experiment. Error bars are standard error of the mean.

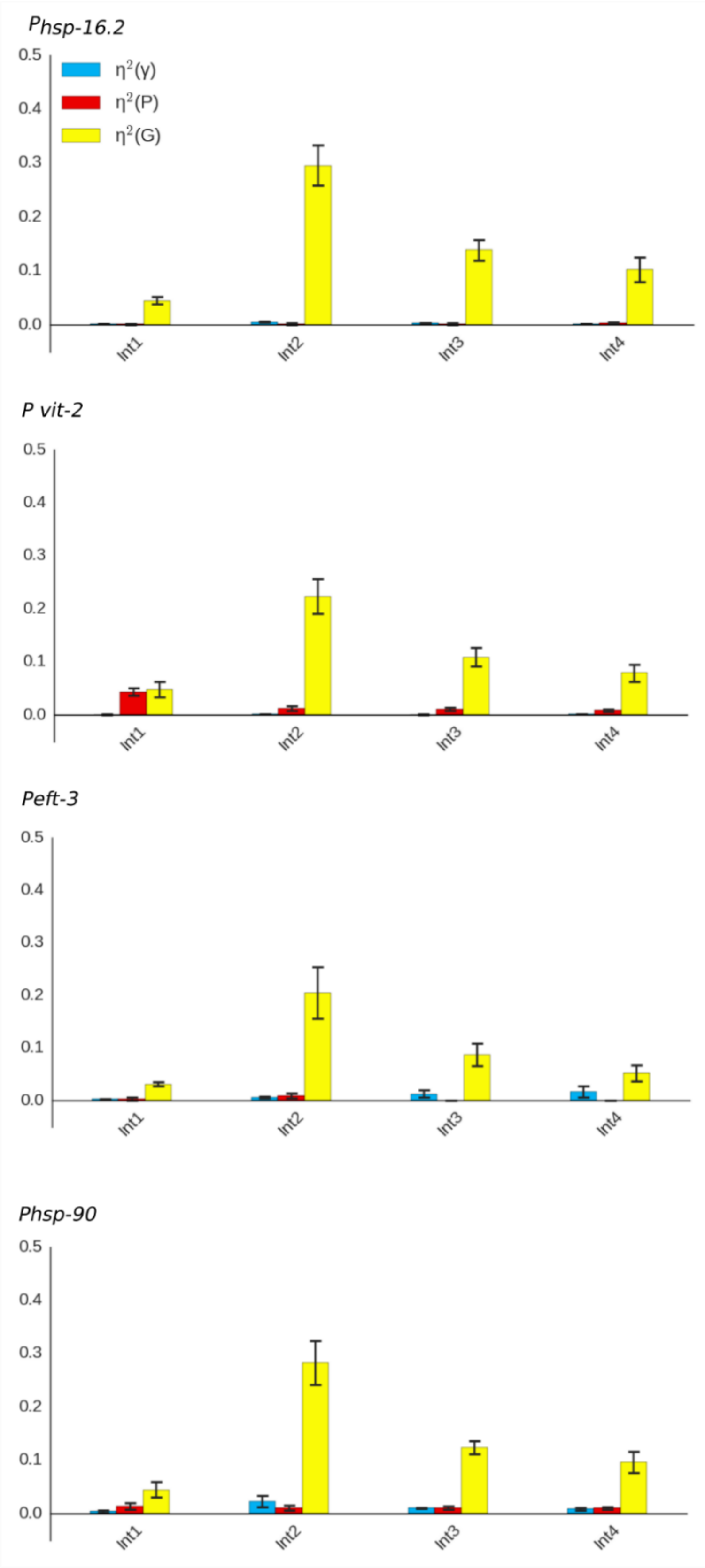

**Supplementary Figure S10. Type II experiments**

**Supplementary bargraphs (no promoter splitting)**

Correlated and uncorrelated variation for different gene pairs in cells in intestine rings 1-4. Uncorrelated variation combines stochastic noise of transcription/translation or variable allele access – $\eta^2(\gamma)$, and variation in pathway activation $\eta^2(P)$. Correlated variation results from variation in gene expression capacity $\eta^2(G)$. These bar graphs are composed of data from three independent experiments quantifying expression from two cells per ring per animal for at least ten animals per experiment. Error bars are standard error of the mean.

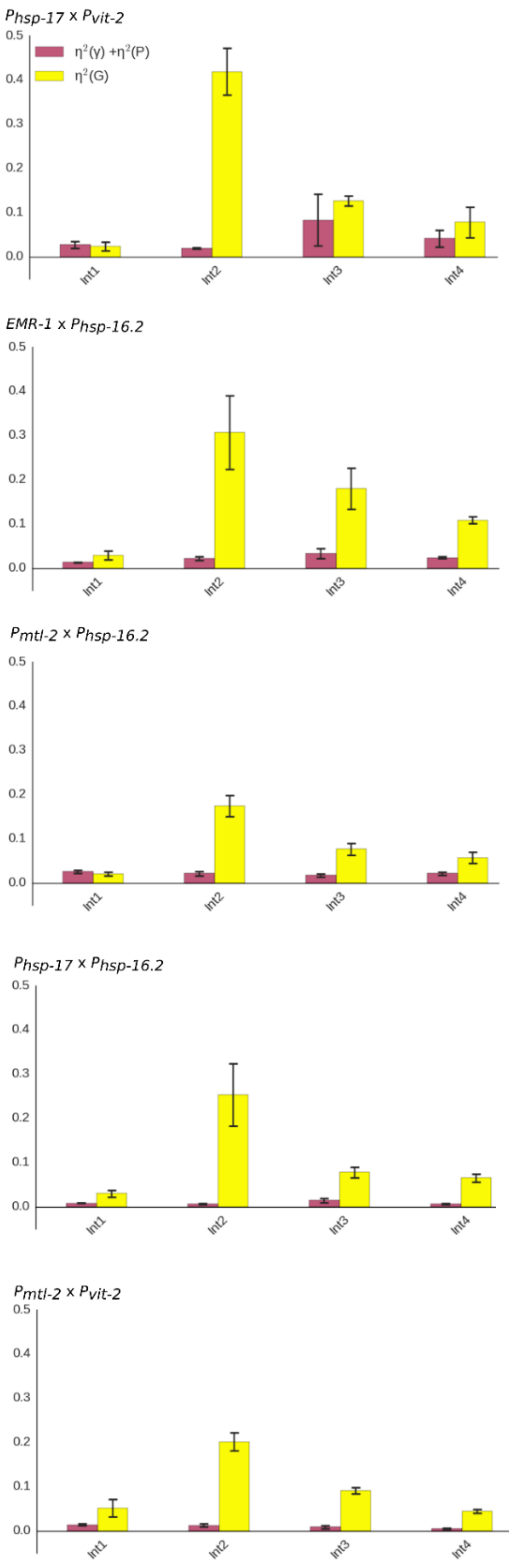

**Supplementary Figure S11. Additional Experiments Quantifying Expression of Nuclear Proteins.** Replicate experiments showing 3-dimensional scatterplots of nuclear fusion proteins Lamin:BFP (single intergenic copy, Chromosome I), His2B:GFP (knockin) and Emerin:mCherry (single intergenic copy, chromosome II), shown in Fig. 4. The data in these scatter plots were generated by measuring individual intestine cell expression levels of all three reporter genes for six to eight cells in each of at least ten animals in two additional independent experiments.

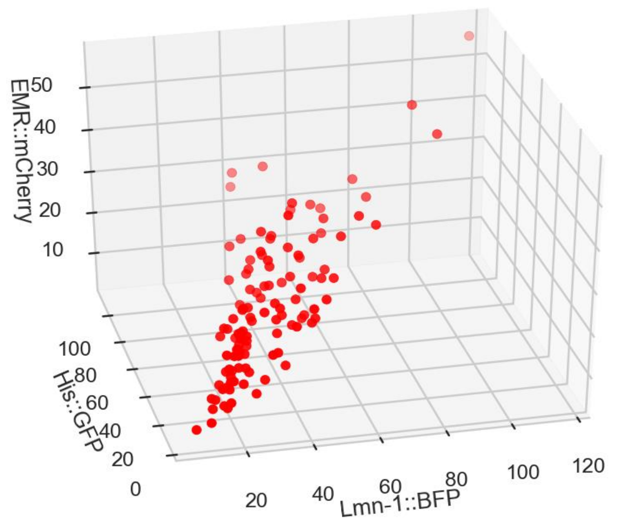


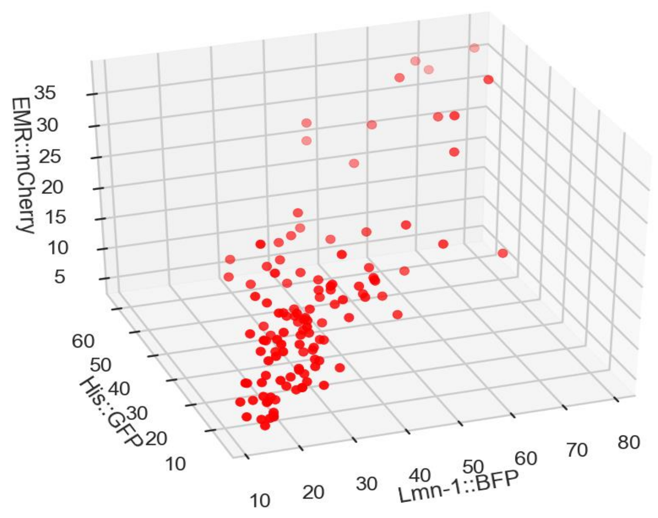

**Supplementary Figure S12. Animals that respond better to heat shock produce and maintain protein better.** **a.** Scatterplot of whole animal average values fluorescent timer protein expressed from *eft-3* promoter ($P_{eft-3}$*::timer*). Linear dependence of young and old protein fractions indicates that bright animals are better at both protein production and maintenance. **b**. Whole animal average abundance of young and old fraction of timer protein with and without heat shock in arbitrary units (tiff counts). The three plots correspond to three independent biological replicates quantifying expression from at least 30 whole animals per group per experiment. Error bars are standard error of the mean.

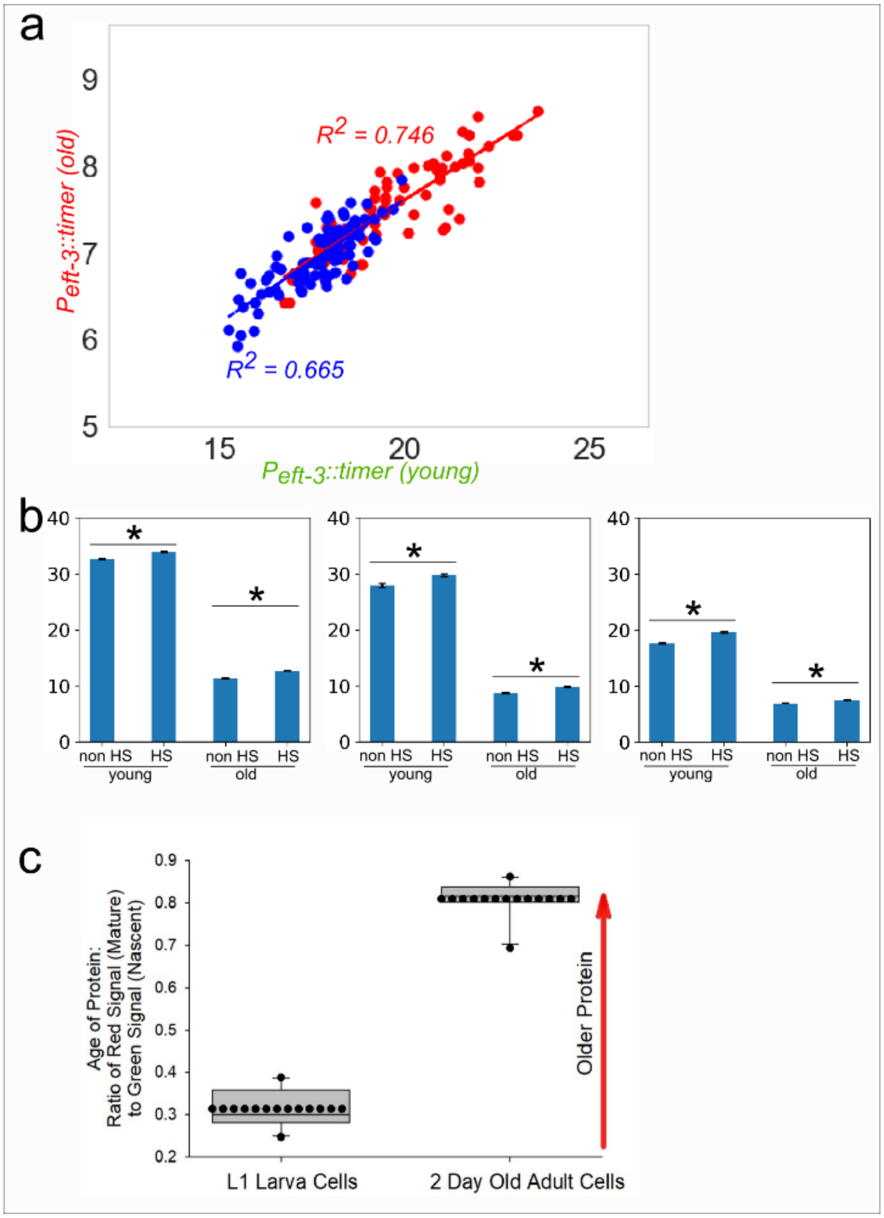


**c**. Timer protein works as expected. To ensure that timer protein reports on average protein age, we examined ratio of old to new timer fraction in approximately ten L1 larvae and in approximately ten, two-day old adults. As expected, slowdown of protein turnover with age increased the ratio of old to new timer protein. The boundary of the box closest to zero indicates the 25th percentile, a line within the box marks the median, a dash within the box marks the average, and the boundary of the box farthest from zero indicates the 75th percentile. Whiskers above and below the box indicate the 90th and 10th percentiles.

**Supplementary Material Section 1: Materials and Methods**

Strains Used, including Creation of Strains Genome Engineering and Creation of *C. elegans* Strains via Mating

We generated strains for these studies using MosSCI transgenesis[1]. We also used some knockins created via CRISPR (his-72::GFP, vit-2::GFP). We also used some standard genetic mutants: CB1370 (*daf-2(e1370)*), CF1038 (*daf-16(mu86)*), NS3099 (*nhr-49(nr2041)*), PS3551 (*hsf-1(sy441)*). We also used some transgenes from prior studies that were multicopy; RBW2 contains a biolistically integrated transgene (*emr-1::gfp)* and was described previously[2]. TJ2741 contains a multicopy version of $P_{hsp\text{-}16.2}$*::gfp*, *zIS2735* [$P_{hsp\text{-}16.2}$*::egfp::*$T_{unc\text{-}54}$] V, which shows the same physiological worm to worm variation as the several other $P_{hsp\text{-}16.2}$ reporters[2]. We used this particular version because it was genetically compatible with the *let-60(n1046)* IV mutation and enabled our sorter to detect signal in the heatshock-pathway-attenuating *let-60(n1046)* background. We made the DNA constructs fusing promoter sequences to the coding sequence of different fluorescent proteins (XFPs) and the terminator of *unc-54* by using yeast gap repair[3]. All reporter constructs carried the 5'UTR (the upstream regulatory sequences including the promoter) of the selected genes (i.e., $P_{hsp\text{-}16.2}$) fused to an XFP coding sequence (*megfp*, *mcherry*, *mtagbfp2*, *mneptune*) and the 3'UTR of *unc-54*. We integrated fluorescent reporters into Chromosome II in the parent strain RBW6699 (an outcrossed version of EG6699 without an extrachromosomal array; *ttTi5605 II; unc-119(ed9)*), unless otherwise noted. To construct DNA for transgenes, we introduced reporter sequences into a vector that targeted the DNA to be inserted at the chromosome II MosSCI transposon at site *ttTi5605*. We then injected each construct at 50ng/µL into EG6699 animals and recovered single copy insertions that we size and site validated by PCR. Thus, through transformation and/or standard genetic crosses, we created the following strains:

RBW2661 (*hutSi2661[*$P_{hsp\text{-}90}$*::megfp::*$T_{unc\text{-}54}$ *+ Cbr-unc-119(+)]* II), RBW2642 (*hutSi2642[*$P_{hsp\text{-}90}$*::mcherry::*$T_{unc\text{-}54}$ *+ Cbr-unc-119(+)]* II), RBW2601 (*hutSi2601[*$P_{hsp\text{-}16.2}$*::megfp::*$T_{unc\text{-}54}$ *+ Cbr-unc-119(+)]* II), RBW2561 (*hutSi2561[*$P_{hsp\text{-}16.2}$*::mcherry::*$T_{unc\text{-}54}$ *+ Cbr-unc-119(+)]* II), RBW2621 (*hutSi2621[*$P_{vit\text{-}2}$*::megfp::*$T_{unc\text{-}54}$ *+ Cbr-unc-119(+)]* II), RBW2581 (*hutSi2581[*$P_{vit\text{-}2}$*::mCherry::*$T_{unc\text{-}54}$ *+ Cbr-unc-119(+)]* II), RBW2531 (*hutSi2531[*$P_{mtl\text{-}2}$*::mcherry::*$T_{unc\text{-}54}$ *+ Cbr-unc-119(+)]* II), ARM1 (*let-60(n1046)*) IV; *jjls699[*$P_{lmn\text{-}1}$*::emr-1::gfp::*$T_{unc\text{-}54}$ *+ unc-119(+)]* ?, not II), ARM6 (*wamSi6[*$P_{eft\text{-}3}$*::mtagbfp2::*$T_{unc\text{-}54}$ *+ Cbr-unc-119(+)]* II), ARM5 (*wamSi5[*$P_{eft\text{-}3p}$*::mneptune::*$T_{unc\text{-}54}$

+ *Cbr-unc-119(+)]* II), RBW3211 (*hutSi3211[*$P_{hsp-17}$*::megfp:*$T_{unc-54}$ + *Cbr-unc-119(+)]* II), TJ2741 ((*zIs2735[*$P_{hsp-16.2}$*::gfp::*$T_{unc-54}$*]*) V; *let-60(n1046)* IV), RBW99 (*oxSi259[*$P_{eft-3}$*::megfp::*$T_{unc-54}$ + *Cbr-unc-119(+)]* I; *oxTi179[*$P_{rps-27}$*::*$Neo^R$*::*$T_{unc-54}$ + *unc-18(+)]* II); RBW2 (*zSi3002[*$P_{hsp-16.2}$*::mcherry::*$T_{unc-54}$ + *Cbr-unc-119(+)*] II; *jjls699[*$P_{lmn-1}$*::emr-1::gfp::*$T_{unc-54}$ + *unc-119(+)]* ?), ARM179 is the cross of NB147 ([emr-1::mCherry + unc-119(+)])II, ARM164 ([lmn-1::mtagBFP2 + unc-119(+)) I and LP148 ([his-72::gfp + LoxP unc-119(+) LoxP]) III), BCN9071 ([vit-2::gfp]), ARM183 is a stable cross of BCN9071 and RBW2642, ARM180 (Peft-3::timer), ARM215 Is a stable cross of RBW6173 ($P_{eft-3}$::GFP) and RBW2581 ($P_{vit-2}$::mCherry).

| Table S4: List of Crosses to make different F1 animals for Type I & II Exps |
|---|
| $P_{daf-21}$*::gfp* (RBW2661) x $P_{daf-21}$*::mcherry* (RBW2642) |
| $P_{daf-21}$*::gfp* (RBW2661) x $P_{vit-2}$*::mcherry* (RBW2581) |
| $P_{hsp-16.2}$*::gfp* (RBW2601) x $P_{daf-21}$*::mcherry* (RBW2642) |
| $P_{hsp-16.2}$*::gfp* (RBW2601) x $P_{hsp-16.2}$*::mcherry* (RBW2561) |
| $P_{hsp-16.2}$*::gfp* (RBW2601) x $P_{vit-2}$*::mcherry* (RBW2581) |
| $P_{vit-2}$*::gfp* (RBW2621) x $P_{vit-2}$*::mcherry* (RBW2581) |
| $P_{mtl-2}$*::mcherry* (RBW2531) x $P_{daf-21}$*::gfp* (RBW2661) |
| $P_{mtl-2}$*::mcherry* (RBW2531) x $P_{hsp-16.2}$*::gfp* (RBW2601) |
| $P_{mtl-2}$*::mcherry* (RBW2531) x $P_{vit-2}$*::gfp* (RBW2621) |
| $P_{eft-3}$*::mtagbfp2* (ARM6) x $P_{daf-21}$*::mcherry* (RBW2642) |
| $P_{eft-3}$*::mtagbfp2* (ARM6) x $P_{hsp-16.2}$*::mcherry* (RBW2561) |
| $P_{eft-3}$*::mtagbfp2* (ARM6) x $P_{mtl-2}$*::mcherry* (RBW2531) |
| $P_{eft-3}$*::mtagbfp2* (ARM6) x $P_{mtl-2}$*::mcherry* (RBW2531) |
| $P_{eft-3}$*::mtagbfp2* (ARM6) x $P_{eft-3}$*::mneptune* (ARM5) |
| $P_{hsp-17}$*::gfp* (RBW3211) x $P_{daf-21}$*::mcherry* (RBW2642) |
| $P_{hsp-17}$*::gfp* (RBW3211) x $P_{hsp-16.2}$*::mcherry* (RBW2561) |
| $P_{hsp-17}$*::gfp* (RBW3211) x $P_{mtl-2}$*::mcherry* (RBW2531) |
| $P_{hsp-17}$*::gfp* (RBW3211) x $P_{vit-2}$*::mcherry* (RBW2581) |
| $P_{hsp-17}$*::gfp* (RBW3211) x $P_{eft-3}$*::mneptune* (ARM5) |
| $P_{hsp-16.2}$*::mcherry* (RBW2561) x *emr-1::gfp* (stable RBW2) |

Culture conditions

We maintained animals on NGM plates seeded with live OP50 *E. coli* at 20°[4]. We ensured that the stocks were not starved for at least two generations prior to use for experimentation. Single-color reporter animals (see above) were mated to produce two-color F1 progeny for microscopy analysis. As a result we analyzed expression of reporters driven by identical or different promoters integrated at the identical loci on both copies of chromosomes II. The only exception is RBW2 strain that carried a transcriptional reporter ($P_{hsp-16.2}$*::mCherry*) and *emr-1::gfp* fusion transgene on different chromosomes and was maintained as a stable strain. Animals that carried $P_{hsp-16.2}$ – driven reporters were heat shocked on the first day of adulthood by exposure to 35°C for one hour on a solid NGM plate. All reporter strains were analyzed on the second day of adulthood. We analyzed crosses listed in Table S4.

Mounting animals for microscopy

We prepared animals for microscopy as previously described in[2]. Briefly, we prepared a 1% agarose pad (about 1-2 $cm^2$) on a standard glass microscope slide. We used worm anesthesia solution containing 0.2% tricaine and 0.02% tetramisole (Sigma, Inc., St. Louis) in M9. We carefully picked animals off their NGM plates and into the drop of anesthesia on the pad. We allowed the worms to swim off the pick into the drop. After that, we covered animals with a glass coverslip. We waited until worms ceased most active movements (about 15-25 minutes), and placed the prepared slide on the slide holder mounted on the microscope stage.

Microscopic image acquisition

We acquired images of animals expressing multiple fluorescent proteins in a similar fashion to Mendenhall et al, 2015[2]. Briefly, we used a 40X 1.2 NA water immersion objective and acquired a z stack of images of each animal in one field of view. Thus, we were able to capture intestine cells in rings one through four of the intestine. We used minimal laser power to prevent photobleaching and heat damage to the cells. We adjusted gain and laser power to ensure that the reporter signals fit within the dynamic range of the photomultiplier tubes and did not saturate them. We only imaged animals lying on their left side to prevent signal loss attributable to imaging through the gonad. We digitally rotated the field of view to arrange the animals into diagonal orientation to fit maximal number of intestine cells in a field of view. We then acquired sequential z-slices to span most of the intestine cells' depth. We acquired an optical slice every two micrometers from the proximal starting point at the objective-proximal side of the intestine, and continued imaging until we acquired signal from intestine cell nuclei in the first four rings. We set optical slice thickness to two micrometers (all photons from plus or minus two micrometers of the image plane). Imaging of each animal typically took two minutes at 1024x1024 resolution. Each pixel value was the average of four samples. We imaged approximately ten animals per experiment.

Image Cytometry

We first determined the orientation of the animals in images and then identified individual cells as described previously, in Mendenhall et al, 2015[2]. As the intestine is anchored anteriorly and posteriorly we used the known positions of the anteriormost cells to determine the identity of adjacent cells. Once we identified a cell, we measured signal within an equatorial slice of the cell's nucleus, as a proxy for the whole cell, as previously described; the nuclear signal is nearly perfectly correlated with the cytoplasmic contents. We used the ImageJ software as well as custom built Nuclear Quantification Support Plugin for nucleus segmentation and signal quantification, we call C Entmoot (Alexander Seewald, Seewald Solutions, Inc., Vienna). The algorithm uses a meeting of decision trees (an Ent Moot – a meeting of tree-beings; from J. R. R. Tolkien's *The Lord of the Rings*) used to delineate a nuclear boundary based on the drop in signal intensity, trained on user delineated images (Seewald et al, manuscript in preparation). The values of the nuclei from binucleate cells were averaged.

Sorting Animals in Flow

For an extensive description of our flow cytometry methods, please see Mendenhall et al, 2015[2]. Also bear in mind that these experiments require the growth of large amounts of animals for sorting – more so than for simply measuring the distribution of a population. Briefly, for the animals bearing the *let-60* mutation to be sorted on $P_{hsp\text{-}16.2}$*::EGFP* expression level, we grew a hypochlorite-synchronized population of animals to gravid adulthood, as in [2,5]. For animals we would sort during the L1 diapause on $P_{eft\text{-}3}$*::GFP* expression level, we performed a hypochlorite synchronization and allowed the animals to hatch and enter the L1 diapause in a 3-5mm deep pool of S-basal swirling at approximately 30 rpms on an unseeded 10cm NGM petri dish. At the start of worm flow sorting, we first flowed a sample of the population of animals to determine the distribution of values in order to select the animals at the extremes of the distribution of fluorescent reporter gene signal; we measured between

2-300 animals to get an estimate of the distribution of the population. We then selected animals at the top or bottom 5-10%, or selected all animals, and sorted between 50-250 animals per group (bright dim, unselected). Actual numbers of animals examined were often less due to loss of animals from escape onto the side of the plate. We hid the identity of each group from the person scoring to prevent bias in scoring. Scoring proceeded immediately after sorting for $P_{hsp\text{-}16.2}$*::EGFP* expression levels for adult animals bearing the *let-60(n1046)* mutation. To score, we counted the number of distinct growths emanating from the ventral hypodermis. A distinct growth was considered a protrusion from the soma unconnected to any other apparent protrusion; we verified that these protrusions were caused by the growth of extra cells using strain ARM1, shown in Supplementary Figure 7. The *let-60* gain of function mutation decreases the expression level of the heatshock response, necessitating the use of the multicopy insertion to ensure detection of GFP signal in this genetic background. We showed previously that the single copy and multicopy reporters have identical lifespan prediction capabilities and the same amount of worm-to-worm variation in biomarker expression. For larvae expressing $Neo^R$, we used a high magnification stereoscope and scored animals for development to gravid adulthood (at least one fertilized embryo in the uterus) after 96 hours of development on food, after being sorted on $P_{eft\text{-}3}$*::GFP* expression while in the L1 diapause (72 hours for experiments on regular NGM).

Whole Animal and Whole Intestine Slide-based Epifluorescent Image Cytometry.

For whole animal/intestine analysis animals were anesthetized with 0.2%Tricaine/0.02%Tetramisole and then mounted on 1% agarose gel pads and covered with cover slips. Animals were imaged on Leica Fluorescent Dissecting scope using MicroManager program for image acquisition. For whole intestine analysis of fluorescence in Pvit-2xPhsp-16.2 and Pvit-2xPeft-3 animals we segmented intestines with thresholding in ImageJ using Pvit-2 fluorescent channel and measured average signals from both reporters in the segmented area. For whole body analysis of fluorescence in Peft-3::timer animals we segmented entire animals with thresholding in ImageJ and measured average signals in each channel in the segmented area.

Correlations Between Embryos, Larva and Adults

For correlations between embryos and adults or between larva and adults, we compared average signal intensities from the cells measured. To measure vit-2::GFP in embryos, we isolated 2-cell embryos and focused on the equatorial plane of the two nuclei, captured an image, and then got the average voxel value for the whole embryo. We did the same thing for whole L1 larvae without anesthesia, focusing on the equatorial plane of the pharynges of animals. We compared larval or embryonic values to the average of adult animals' intestine cells by averaging cell voxel intensity values from the adult cells we measured.

Lifespan and Fecundity After UV Irradiation

We placed populations of approximately 100, age-synchronized young adult animals (approximately 60 hours development on NGM with OP50 post starved L1 diapause) in a StrataLinker and irradiated them with 1000 Joules of ultraviolet radiation. We took 50 animals from each group to conduct lifespans and we took 5 animals from each group to perform fecundity assays. We also performed self-fertility fecundity assays on wild-type animals that had never been heat shocked.

Statistical Analysis

We used Sigma Stat (Systat Software, Inc., San Jose) for statistical analyses of the bright and dim animals we sorted. We first determined if measurements comprising each dataset were normally distributed, then, depending on the results of those tests, used appropriate parametric or non-parametric statistics to determine if there was significant difference in any measured parameters. Details of specific tests are shown in figure legends. Additional details regarding grouping of different types of measurements for calculations of $\eta^2$(G, P, or γ) and statistical analyses of different variation bins see Supplementary Materials Section 2: Analytical Framework. For all reported coefficients of correlation or determination, we used a Pearson correlation.

### Supplementary Material Section 2: Analytical Framework

#### S2.1 Analytical Framework: Defining Pathway Output and Expression Capacity

For quantitative analysis we adapted the analytical framework developed previously by Coleman-Lerner et al and briefly described below. We considered the production of fluorescent proteins (e.g. GFP and mCherry) in individual cells to be a measure of activities of the promoters that regulate their expression. Furthermore, to understand cell-to-cell and animal-to-animal differences in activities of various promoters we considered the amount of fluorescent protein produced to be the product of two subsystems: "pathway" and "expression".

"Pathways" is the first subsystem. For instance, for a heat shock inducible promoter $P_{hsp\text{-}16.2}$, the input to pathway is the activation of transcription factors, e.g. HSF-1, that bind to *hsp-16.2* promoter, and the output is the activation of the inducible promoter that drives the fluorescent protein. Pathway output depends on the summed activity of upstream, DNA-bound transcription factors.

For a given cell *i,* pathway output is proportional to $P_i$, the time-averaged of the level of pathway activation. We separate $P_i$ as $P_i = L_i + \lambda_i$ where $L_i$ is the expectation value of $P_i$ for a given cell type in a cohort of isogenic age-matched animals, and $\lambda_i$ is the stochastic fluctuation term for that cell. We refer to $L_i$ as the pathway power. The pathway "power" is a function of the activities of upstream signaling molecules and transcription factors that lead to activation of the promoter. We refer to cell-to-cell differences in $P_i$ as "variation in pathway power", and we describe differences that result from the stochastic fluctuation term $\lambda_i$ as "transmission noise". Thus, we can treat the output of the cell as being decomposed into an expectation value, which depends on the number and activity of the molecules that comprise the pathway, and a stochastic fluctuation that occurs because of the inherent randomness of the activity of the pathway, as well as its composition over the time of the experiment. Different cells may have different pathway capacities, and therefore, can have different values of $L_i$.

The second subsystem consists of the sequence of events from gene transcription through protein translation. We call this subsystem "*expression*". It includes transcriptional initiation, elongation, mRNA maturation, nuclear export, and mRNA translation and degradation (of protein and mRNA). We use our data to measure cell-to-cell variation in expression, but we cannot isolate the contribution of each part of the expression machinery to the overall variation.

The output of the expression subsystem is the total amount of the fluorescent reporter protein, and the input is proportional to the level of promoter activity (the output of the pathway subsystem). For analytical purposes, we assume here that the expression per unit of input is independent of the level of input. For cell *i,* we describe the expression per unit input as the variable $E_i$, where $E_i$ is given by the sum of $G_i$ and $\gamma_i$. The quantity $G_i$ is the expectation value of the expression per unit input for cell *i,* and this quantity may vary from cell to cell. We refer to $G_i$ as the “expression capacity” for cell *i*, and we refer to the cell-to-cell differences in $G_i$ as “variation in expression capacity”. The quantity $\gamma_i$ is the stochastic fluctuation that occurred in cell *i*, and we refer to differences that result from this stochastic fluctuation term as “expression noise.”

Using this model, we described the total amount of GFP, $y_i$, in cell *i* as

$$y_i = (L_i + \lambda_i) \text{ x } (G_i + \gamma_i) \text{ x } \Delta\text{T} \quad (1)$$

where ΔT is the time of reporter protein production. For heat inducible promoter we can consider ΔT as the time since the heat shock. For constitutive promoters like *vit-2* we cannot precisely measure what fraction of the observed reporter protein pool was produced in the same time frame. Thus, we started this work under “steady state” assumption that reporter protein synthesis and degradation are balanced in young adult animals. In that case reporter proteins level directly mirror activity of the promoters and expression capacity. Highly correlated expression of the examined promoters suggests that this assumption is applicable and acceptable.

**S2.2 Extracting Pathway and Expression Capacity Information from the Data**

**Variance and Covariance**

We define the variance of a given quantity *x* for a population of cells as

$$\sigma^2 = \frac{1}{N}\sum(x_i - \bar{x})^2 \quad (2)$$

$$= \overline{(x_\iota - \bar{x})} \quad (3)$$

where *N* is the number of cells in the population. Since, we examined homologous cells of *C. elegans* nematodes, this number is equal to the number of animals in the population. We use the overbar symbol ($\bar{x}$) to represent the population average. Similarly we define the covariance, *Cov*(*x*,*y*), of two quantities *x* and *y* as

$$Cov(x, y) = \frac{1}{N}\sum(x_i - \bar{x})(y_i - \bar{y}) \quad (4)$$

$$= \overline{(x_\iota - \bar{x})(y_\iota - \bar{y})} \quad (5)$$

The covariance will be non-zero if x and y are correlated (or anti-correlated).

The correlation coefficient, ρ(x,y), is the covariance scaled by the standard deviations. Specifically

$$\rho(x,y) = \frac{Cov(x,y)}{\sigma(x)\sigma(y)} \quad (6)$$

**S2.3 Calculating Variance and Covariance of** G, γ**, L, and** λ

As discussed above we assumed that in standard conditions, the average pathway output (*P*) for a given cell has a pathway power *L* associated with it, along with a stochastic term, *λ*. Thus, we have *P=L+λ.* Similarly, the expression subsystem of the cell (*E*) contains a capacity term G and a stochastic term γ.

Thus, the amount of fluorescent reporter protein ($y_i$) produced for a given cell *i* is the product of the terms

$$y_i = (L_i + \lambda_i) \times (G_i + \gamma_i) \text{ x } \Delta\text{T} \quad (7)$$

$$= (L_iG_i + L_i\gamma_i + \lambda_iG_i + \lambda_i\gamma_i) \times \Delta\text{T} \quad (8)$$

Since the expectations of the stochastic fluctuation terms are zero, and since their fluctuations are uncorrelated to other terms, we expect that the population average of $y_i$ reduces to the population average of the quantity $L_iG_i\Delta T$.

To calculate the average of y, we re-write the product $L_i$ x $G_i$ in terms of population averages and deviations from the average. For these two terms, we use a capital delta (*Δ*) to represent a deviation from a population average. We get

$$L_i = \bar{L} + \Delta L_i \quad (9)$$

$$G_i = \bar{G} + \Delta G_i \quad (10)$$

We then calculate the population average of *y* to be

$$\bar{y} = \frac{\Delta}{T}\sum(L_i \times G_i) \tag{11}$$

$$= (\bar{L} \times \bar{G} + Cov(L,G))\Delta T \tag{12}$$

where we used equation (4) for the definition of covariance and the fact that *ΔG=ΔL=0*. Thus the average number of GFP molecules is the product of *L* and G plus an extra term to account for their correlation.

To calculate the variance on *y*, we re-write equation (7) in terms of equations (9) and (10). We get

$$y_i = (\bar{L} + \Delta L_i + \lambda_i)\Delta T \times (\bar{G} + \Delta G_i + \gamma_i) = \tag{13}$$

$$= \left(1 \frac{\Delta L_i}{\bar{L}} + \frac{\lambda_i}{\bar{L}}\right) \times \left(1 + \frac{\Delta G_i}{\bar{G}} + \frac{\gamma_i}{\bar{G}}\right) \times \bar{G} \times \bar{L} \times \Delta T \tag{14}$$

$$= \left(1 + \frac{\Delta L_i}{\bar{L}} + \frac{\lambda_i}{\bar{L}}\right) \times \left(1 + \frac{\Delta G_i}{\bar{G}} + \frac{\gamma_i}{\bar{G}}\right) \times \bar{y}(1 - \frac{Cov(L,G)\Delta T}{\bar{y}}) \tag{15}$$

$$\approx y \times (1 + \frac{\Delta L_i}{\bar{L}} + \frac{\lambda_i}{\bar{L}} + \frac{\Delta G_i}{\bar{G}} + \frac{\gamma_i}{\bar{G}}) \tag{16}$$

where we used equation (12) to make the substitution for $\bar{L} \times \bar{G} \times \Delta T$ in going from equation (14) to (15); and where we have dropped, in going from equation (15) to (16) all higher order terms of fractional deviations from the means. These higher order terms are various products of $\frac{\Delta L_i}{\bar{L}}$, $\frac{\lambda_i}{\bar{L}}$ , $\frac{\Delta G_i}{\bar{G}}$ , and $\frac{\gamma_i}{\bar{G}}$ . We assume that each of the higher order terms is small relative to the lower order terms that are retained in equation (16), and therefore we have neglected them. The term $\frac{Cov(L,G)\Delta T}{\bar{y}}$ is second order, and has also been neglected. Discussion of the magnitude of the error introduced by this approximation is provided in Coleman-Lerner et al.

We use equation (16) to calculate the variance on the number of GFP molecules using the definitions of variance and covariance above. We get

$$\frac{\sigma^2(y)}{\bar{y}^2} = \frac{\sigma^2(L)}{\bar{L}^2} + \frac{\sigma^2(\lambda)}{\bar{L}^2} + \frac{\sigma^2(G)}{\bar{G}^2} + \frac{\sigma^2(\gamma)}{\bar{G}^2} + 2\rho(L,G)\frac{\sigma(L)}{\bar{L}}\frac{\sigma(G)}{\bar{G}} \tag{17}$$

where $\rho(L,G)$ is the correlation coefficient between $L$ and G.

The correlation coefficient is always between -1 and 1, and a value of 0 corresponds to no correlation. There may be no correlation between $L$ and G, but, for example, one possibility that would lead to a correlation is that cells with higher expression capacity may have stronger or weaker pathway output, perhaps because they have higher amounts of positive regulators or negative regulators of the pathway, respectively. This would show up as a positive or negative value for $\rho(L,G)$, respectively.

We do not expect that the stochastic fluctuations λ or γ are correlated to any other variables, and thus the terms $\rho(L,\lambda)$, $\rho(L,\gamma)$, $\rho(\gamma,G)$, $\rho(\lambda,G)$, *and* $\rho(\lambda,\gamma)$ were omitted from equation (17).

All the variances in equation (17) are expressed as fractions of the means squared. We define the variables

$$\eta(L) \equiv \sigma(L)/\,\bar{L} \qquad (18)$$

$$\eta(\lambda) \equiv \sigma(\lambda)/\,\bar{L} \qquad (19)$$

$$\eta(G) \equiv \sigma(G)/\,\bar{G} \qquad (20)$$

$$\eta(\gamma) \equiv \sigma(\gamma)/\,\bar{G} \qquad (21)$$

The quantity $\eta$ is simply the width of different distributions expressed as a fraction of their means. For the stochastic variables $\lambda$ and $\gamma$, we will refer to $\eta(\lambda)$ and $\eta(\gamma)$ as the "noise" associated with those quantities, and for the non-stochastic variables $L$ and G, we will refer to $\eta(L)$ and $\eta(G)$ as the "variation."

We quantified cell-to-cell variation in system output using "normalized variance" ($\eta^2 = \sigma^2/\mu^2$) rather than "noise strength" ($\sigma^2/\mu$), a measure others have used to describe deviations from purely stochastic Poisson-type biological processes. We used normalized variance for a number of reasons. First, we found that most of the cell-to-cell differences in system behavior we reported are not due to stochastic differences in signal transmission or gene expression, as described in the main text. Second, use of $\eta^2$ allowed examination of different amounts of variation in terms of the fraction of the mean. Third, and most important, because $\eta^2$ is unitless, it allowed direct comparison of different measurements, for example, from different fluorescent proteins, and it allowed the definition of total variation as the sum of individual sources of variation plus additional terms to account for correlations.

We re-write equation (17) in terms of η as

$$\eta^2(y) = \eta^2(P) + \eta^2(G) + \eta^2(\gamma) + 2\rho(L,G)\eta(L)\eta(G) \quad (22)$$

We have written $\eta^2(P)$ for $\eta^2(L)+\eta^2(\lambda)$ since the data will not be able to distinguish the cell-to-cell variation in pathway output from the noise or stochastic fluctuations. The term $\eta^2(P)$ is the variation in average pathway output per unit time, which, because pathway output is given by $P\Delta T$ and $\Delta T$ is the same for every cell, is identical to variation in pathway output.

To separate $\eta(G)$ from $\eta(P)$ and $\eta(\gamma)$ in the data from the two-promoter, two-color experiments, discussed further below, we introduce the quantity

$$Z(GFP, mCherry) = \frac{\eta^2(GFP)+\eta^2(mCherry)}{2} - \eta(GFP)\eta(mCherry)\rho(GFP, mCherry)$$

$$= \frac{\eta^2(GFP)+\eta^2(mCherry)}{2} - \frac{Cov(GFP, mCherry)}{\overline{GFP} \times \overline{mCherry}} \quad (23)$$

The quantity $Z$ is the average variance divided by the mean square of the two fluorescence signals GFP and mCherry with the correlated part subtracted out. Thus, $Z$ is a measure of the uncorrelated part of the GFP vs mCherry scatter plot. Depending of the type of the experiment (two identical promoters or two different promoters) $Z$ encompasses distinct sources of variation.

#### S2.4 Two Color Variants Driven by the Same Promoter and Gene Expression Noise

We calculate the correlation coefficient between GFP and mCherry reporter proteins for the case that both genes are expressed in the same cell and with the same promoter as

$$\rho(GFP, mCherry) \equiv \frac{1}{\sigma(GFP)\sigma(mCherry)}\frac{1}{N}\sum_i \Delta(GFP_i)\Delta(mCherry_i) \quad (24)$$

$$= \frac{1}{\eta(GFP)\eta(mCherry)}\frac{1}{N}\sum_i\left(\frac{\Delta L_i}{\bar{L}} + \frac{\lambda_i}{\bar{L}} + \frac{\Delta G_i}{\bar{G}} + \frac{\gamma_{GFP,i}}{\bar{G}}\right)\left(\frac{\Delta L_i}{\bar{L}} + \frac{\lambda_i}{\bar{L}} + \frac{\Delta G_i}{\bar{G}} + \frac{\gamma_{mCherry,i}}{\bar{G}}\right) \quad (25)$$

where we have used equation (16) for the deviations from the mean, $\Delta(GFP_i)$, and an analogous equation for $\Delta(mCherry_i)$. Since the GFP gene has the same promoter as mCherry, $\Delta L_i$ is the same for both color variants. The quantity $\lambda_i$ is the same since stochastic fluctuations in pathway output occur upstream of the promoters (with the exception of the binding of the transcription factors to each individual copy of the reporter genes, see the end of this section), and the quantity $\Delta G_i$ is the same since the proteins are being expressed in the same cell. The stochastic fluctuations in gene expression, $\gamma_i$, however, are different for the two color variants and we write them as $\gamma_{GFP,i}$ and $\gamma_{mCherry,i}$ for GFP and mCherry respectively.

The uncorrelated terms drop out of equation (25) when we perform the population average. We do not expect the stochastic fluctuations to be correlated with any other terms, and equation (25) becomes

$$\rho(GFP, mCherry) = \frac{\eta^2(P)+\eta^2(G)+2\rho(L,G)\eta(L)\eta(G)}{\eta(GFP)\,\eta(mCherry)} \quad (26)$$

Where $\eta^2(P) = \eta^2(L) + \eta^2(\lambda)$ as discussed above.

The quantity $Z(GFP, mCherry)$ as defined in equation (23), gives the contribution of the uncorrelated part of the expression of the two color variants to the average $\eta^2$ of the two colors. Only the gene expression noise is uncorrelated, and we get

$$Z(GFP, mCherry) = \frac{\eta^2(\gamma_{GFP})+\eta^2(\gamma_{mCherry})}{2} \quad (27)$$

$Z(GFP, mCherry)$ is the average of gene expression noise of the two promoters. Since the same promoters are driving the GFP and mCherry genes, we expect equal levels of mRNA for the two variants on average, and therefore the gene expression noise is the same for the GFP and mCherry signal. We get $\eta^2(\gamma_{GFP}) =$

$\eta^2(\gamma_{mCherry}) = \eta^2(\gamma)$ and therefore, from equation (27), $Z(GFP, mCherry) = \eta^2(\gamma)$. Thus, equation (23) gives direct measurement of gene expression noise $\gamma$ when applied to an experiment with two identical promoters:

$$\eta^2(\gamma) = \frac{\eta^2(GFP)+\eta^2(mCherry)}{2} - \frac{Cov(GFP,mCherry)}{\overline{GFP} \times \overline{mCherry}} \qquad (28)$$

<u>We used this equation (28) for practical calculations of gene expression noise of individual promoters.</u>

We note that for the experiment described above, GFP and mCherry were not driven by the same promoter, but rather by identical *copies* of the same promoter. Therefore, stochasticity in the binding of transcription factors to the DNA and in the subsequent activation of transcription contributes to the uncorrelated part of the total variation. In our *ad hoc* model, these molecular steps are part of the pathway subsystem, and therefore fluctuations in this step should contribute to the transmission noise, $\eta^2(\lambda)$. However, due to experimental limitations of the type of experiment we have just described, the noise caused by these molecular steps is, instead, included in the measure of gene expression noise $\eta^2(\gamma)$.

**S2.5 Two Color Variants Driven by Different Promoters and Pathway Variation**

For this case, the calculation of *ρ(GFP,mCherry)* is the same as in equation (25) above except that the pathway activity is different for the two color variants since different promoters are driving the GFP and mCherry genes. If we assume that the two terms *L* and *λ* are uncorrelated for the two promoters then we get

$$\rho(GFP, mCherry) = \frac{\eta^2(G)+\rho(L_{GFP},G)\eta(L_{GFP})\eta(G)+\rho(L_{mCherry},G)\eta(L_{mCherry})\eta(G)}{\eta(GFP)\,\eta(mCherry)} \qquad (29)$$

where we have written $L_{GFP}$ and $L_{mCherry}$ for capacities of the pathways that lead to the activation of the GFP and mCherry gene respectively. The uncorrelated part of the average $\eta^2$ will now include a contribution from the pathway variation; and the quantity *Z*(*gfp*, m*Cherry*) includes this extra contribution. We get

$$Z(GFP, mCherry) = \frac{\eta^2(P_{GFP})+\eta^2(P_{mCherry})}{2} + \frac{\eta^2(\gamma_{GFP})+\eta^2(\gamma_{mCherry})}{2} \quad (30)$$

where $\eta^2(\lambda_{GFP})$ is the transmission noise for the pathway that leads to GFP activation and $\eta^2(P_{GFP}) = \eta^2(L_{GFP}) + \eta^2(\lambda_{GFP})$, and similarly for $\eta^2(\lambda_{mCherry})$ and $\eta^2(P_{mCherry})$.

Thus, when uncorrelated expression variation for two independent promoters includes an average variation in pathways activation and gene expression variations for each promoter. The latter can be calculated from type I experiments. Knowing expression variations $\eta^2(\gamma)$ for each promoter and uncorrelated variation $Z$ from equation (23) allows to calculate of pathway variations $\eta^2(P)$. To split contribution of variation of each pathway we need to perform three experiments with pairwise analysis of three promoters and then solve a system linear equations to calculate $\eta^2(P)$ of each pathway.

Below we give an example with expression of three promoters, *vit-2*, eft-3, daf-21, in cells Int3V and Int3D.

$$Z(P_{eft-3} :: BFP, P_{vit-2} :: mCherry) = \frac{\eta^2(P_{eft-3})+\eta^2(P_{vit-2})}{2} + \frac{\eta^2(\gamma_{P_{eft-3}::BFP})+\eta^2(\gamma_{P_{vit-2}::mCherry})}{2} \quad (31)$$

$$Z(P_{eft-3} :: BFP, P_{hsp-90} :: mCherry) = \frac{\eta^2(P_{eft-3})+\eta^2(P_{hsp-90})}{2} + \frac{\eta^2(\gamma_{P_{eft-3}::BFP})+\eta^2(\gamma_{P_{hsp-90}::mCherry})}{2} \quad (32)$$

$$Z(P_{hsp-90} :: GFP, P_{vit-2} :: mCherry) = \frac{\eta^2(P_{hsp-90})+\eta^2(P_{vit-2})}{2} + \frac{\eta^2(\gamma_{P_{hsp-90}::GFP})+\eta^2(\gamma_{P_{vit-2}::mCherry})}{2} \quad (33)$$

Average gene expression noise of two promoters, e.g. $\frac{\eta^2(\gamma_{P_{eft-3}::BFP})+\eta^2(\gamma_{P_{vit-2}::mCherry})}{2}$ is calculated from type I experiments and equation (28). It is therefore a known value for each cell type and promoter. We calculate quantity $Z$ directly from equation (23). Quantity Z and average expression noise therefore give a particular number of each of the equations above. We can rewrite them in more suitable form to calculate pathways variations

$$\frac{\eta^2(P_{eft-3})+\eta^2(P_{vit-2})}{2} = A_1 \quad (34)$$

$$\frac{\eta^2(P_{eft-3})+\eta^2(P_{hsp-90})}{2} = A_2 \quad (35)$$

$$\frac{\eta^2(P_{hsp-90})+\eta^2(P_{vit-2})}{2} = A_3 \quad (36)$$

Where $A_1$, $A_2$ and $A_3$ are known values. To calculate pathway variation for a particular promoter, e.g. eft-3, we will sum up (34) and (35) and subtract (36):

$$\frac{\eta^2(P_{eft-3})+\eta^2(P_{vit-2})}{2} + \frac{\eta^2(P_{eft-3})+\eta^2(P_{hsp-90})}{2} - \frac{\eta^2(P_{hsp-90})+\eta^2(P_{vit-2})}{2} = A_1 + A_2 - A_3 \quad (37)$$

$$\frac{\eta^2(P_{eft-3})+\eta^2(P_{vit-2})+ \eta^2(P_{eft-3})+\eta^2(P_{hsp-90})-\eta^2(P_{hsp-90})-(P_{vit-2})}{2} = A_1 + A_2 - A_3 \quad (38)$$

$$\frac{2\eta^2(P_{eft-3})}{2} = A_1 + A_2 - A_3 \quad (39)$$

$$\eta^2(P_{eft-3}) = A_1 + A_2 - A_3 \quad (40)$$

Thus, we have got one estimate of $\eta^2(P_{eft-3})$. To obtain it we used results of the experiments involving $P_{eft-3}$, $P_{vit-2}$ and $P_{daf-21}$ reporters in pairwise manner. We say that these 3 pairs of experiments form a 'triangle'. However, we could choose another triangle consisting of experiments with $P_{eft-3}$, $P_{vit-2}$ and $P_{hsp-16.2}$ reporters, or a triangle of $P_{eft-3}$, $P_{hsp-16.2}$ and $P_{hsp-90}$ reporters. To get a better estimate of the true $\eta^2(P_{eft-3})$ value we have calculated it from all three possible triangles and averaged. We did the same for other reporters too. The resulting $\eta^2(P)$ values are shown in Figure 3 bar graphs.

Similarly, we can calculate pathway variation for other promoters for which we have measured their expression noise. In cases when we did not measure expression noise, e.g. *hsp-17* promoter, we did calculate pathway variation for these promoters as well. In these cases we have limited our analysis with comparison of total correlated variation of two promoters to their uncorrelated variation. In all cases we have observed that correlated variation dominates over uncorrelated term:

$$\frac{Cov(P_{hsp-17}::GFP, P_{eft-3}::mNeptune)}{\overline{P_{hsp-17}::GFP} \times \overline{P_{eft-3}::mNeptune}} \gg Z(P_{hsp-17} :: GFP, P_{eft-3} :: mNeptune)$$

**Supplementary Material Section 3: Additional Correlations between Phenotypes and Reporter Genes**

**As in Previous Reports, We Also Find Activities of Gene Products Correlate with the Expression Levels of Fluorescent Reporter Genes at Other Points in Life.** Previous reports showed the *hsp-16.2* and *hsp-90* chaperone biomarkers covaried with the activities of mutant proteins, discerned from the mutant phenotypes that did or did not manifest during larval development. We used the *hsp-16.2* biomarker gene to sort adult animals expressing an incompletely penetrant, dominant Ras gain of function mutation, which causes animals to develop between zero and four hypodermal neoplasias. If animals that had higher Ras activity had higher gene expression capacity that persisted into adulthood, we would expect the *hsp-16.2* biomarker to correlate with the penetrance and expressivity of the Ras mutation. Supplementary Fig. S13 shows that animals that make less of the biomarker have a lower penetrance; brighter animals have more hypodermal neoplasias. This evidence supports the other evidence showing that animals vary in effective gene dosage for multiple different genes simultaneously. Supplementary Fig. S14 shows images of animals in which the nuclear envelope of the somatic cells, including the neoplastic hypodermal cells, is delineated by a green fluorescent protein fusion to Emerin, a nuclear envelope component. Note that, even without seeing the covariation with other genes, the individual animals with distinct numbers of growths emanating from their ventral surface provides obvious evidence of differences in Ras activity between animals. We are only showing evidence of previous Ras activity is covarying with chaperones after adult heat shock. The relationship between elevated Ras and chaperone activity is not new or unknown; chaperone-targeting drugs have been used to suppress Ras activity in cancer treatment[6,7].

Next, we tested the hypothesis that some other, non-native larval gene activity could be predicted by expression of a reporter gene. We tested the hypothesis that we could detect correlations between a ubiquitously expressed reporter that correlated well with chaperone biomarkers, $P_{eft\text{-}3}$*::GFP*, and the activity of the neomycin resistance gene controlled by another promoter, *rps-27*. We sorted diapaused larval (L1) animals on $P_{eft\text{-}3}$*::GFP* expressed from a site on chromosome I; these animals expressed a neomycin resistance gene (NeoR) on a different chromosome (chromosome II) under control of the *rps-27* promoter. If general protein expression capacity (G) was a major mechanism of intestine cell-to-cell (and animal-to-animal) variation, then animals that have more GFP should have more NeoR and resist extreme neomycin concentrations to a greater extent.

Supplementary Fig. S15 shows that we found this to be the case; brighter green animals can grow to adulthood in a high neomycin environment better than dimmer animals.

The data from these experiments is consistent with prior observations of covariation between fluorescent reporters and apparent phenotypes previously reported in larval worms. Taken together with all of the previous data on covariation and the data on covariation between reporter genes in this manuscript, the case for chaperone biomarkers revealing physiological states of different proteome dosage (at least for significant portions of the proteome) seems well supported. For this entire manuscript, the data comprising these covariation measurements consists of many distinctly regulated promoters, proteins and loci (see Table S4), using four distinct fluorescent proteins, originating from the zooxanthellae endosymbionts of three different cnidarians. Thus, these results cannot be an artifact of some property of any single promoter, protein, locus or fluorescent protein (e.g., folding, maturation, quantum efficiency, bleaching, pKa).

**Supplementary Figure 13 Penetrance and Expressivity of the Ras Gain of Function Mutation is Correlated with *hsp-16.2* Expression Levels.** Penetrance of Ras/let-60 gain of function mutation *let-60(n1046)* correlates with the ability to express $P_{hsp\text{-}16.2}$*::GFP* reporter. Chi Square for distributions of bright or dim vs unselected $P < 0.0001$. Difference in penetrance between bright and dim $P < 0.05$, Paired t-test. See Tables S1a-c for numerical data.

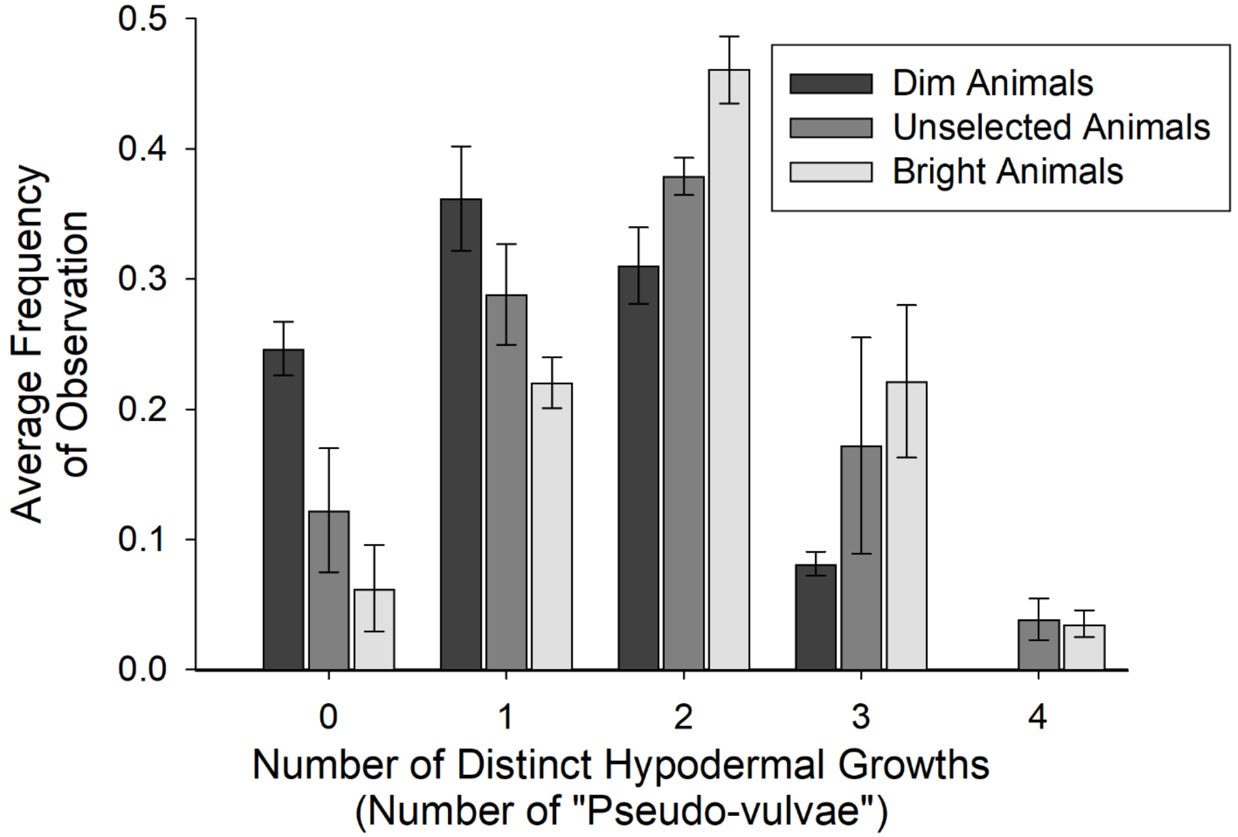

**Supplementary Figure 14. Penetrance and Expressivity of Ras Gain of Function Mutation in *C. elegans.***

Animals express EMR-1::GFP to mark cell nuclei. Wild type and *let-60(n1046)* animals are shown in the top two panels. Some *let-60* mutants do not develop extra pseudo-vulvae/neoplasias (impenetrant, phenotypically wild type). Other *let-60* animals exhibit variable numbers of hypodermal neoplasias. See methods for details of microscopy.

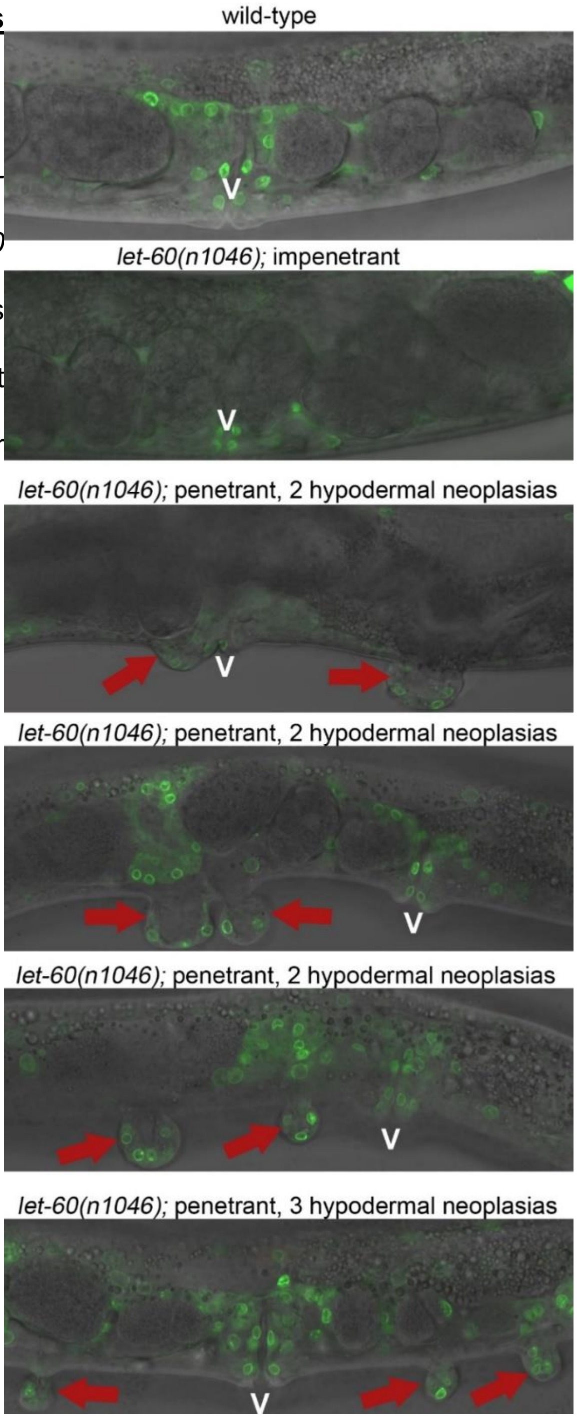

**Supplementary Figure 15. Animals Expressing the NeoR gene that Express more Peft-3::mEGFP in the L1 diapause Develop to Adulthood Better on Neomycin.** Drug resistance conferred by $Neo^R$ expressed from $P_{rps\text{-}27}$ promoter correlates with expression of $P_{eft\text{-}3}$*::EGFP* reporter. $P_{eft\text{-}3}$*::EGFP* reporter does not predict ability to grow in the absence of neomycin (striped bars). High $P_{eft\text{-}3}$*::EGFP* expression predicts a significant difference in the ability to grow to adulthood on a high neomycin concentration (solid bars; $P < 0.05$, Paired t-test). See Tables S2 and S3 for numerical data.

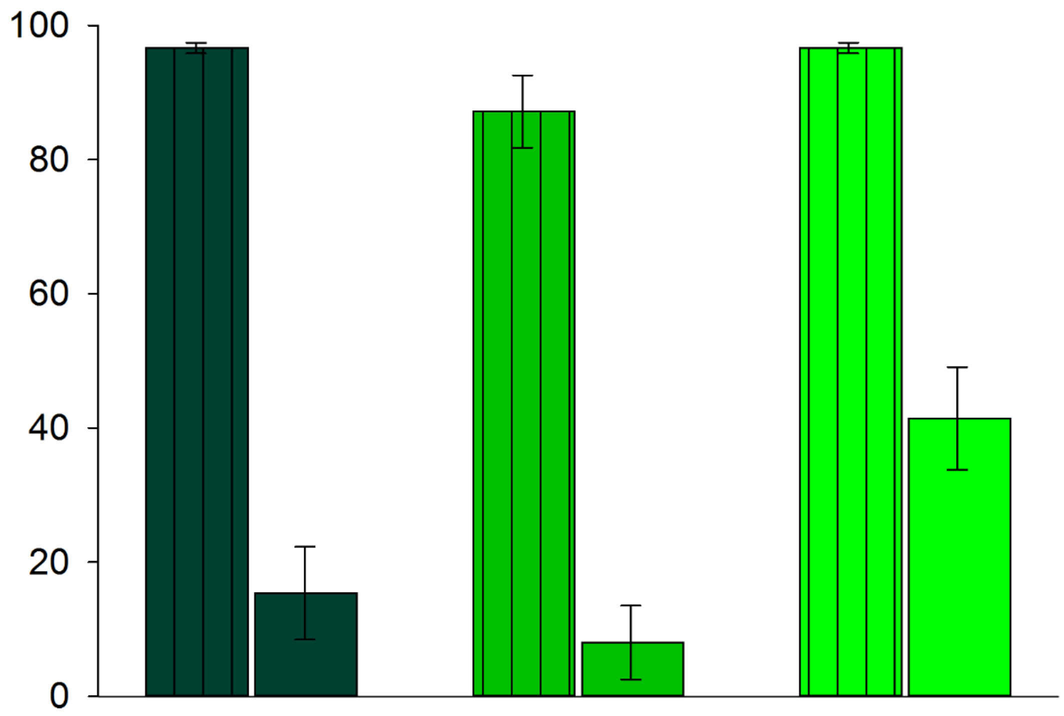

Table S1a Number of hypodermal neoplasias

Experiment 1

| # Growths | Dim | | Unselected | | Bright | |
|---|---|---|---|---|---|---|
| | # Animals | % Animals | # Animals | % Animals | # Animals | % Animals |
| 0 | 10 | 23% | 2 | 3% | 0 | 0% |
| 1 | 19 | 44% | 15 | 21% | 7 | 19% |
| 2 | 11 | 26% | 25 | 35% | 15 | 42% |
| 3 | 3 | 7% | 24 | 34% | 12 | 33% |
| 4 | 0 | 0% | 5 | 7% | 2 | 6% |
| Total # Animals | 43 | | 71 | | 36 | |

Table S1b Number of hypodermal neoplasias

Experiment 2

| # Growths | Dim | | Unselected | | Bright | |
|---|---|---|---|---|---|---|
| | # Animals | % Animals | # Animals | % Animals | # Animals | % Animals |
| 0 | 27 | 29% | 31 | 16% | 6 | 7% |
| 1 | 30 | 32% | 66 | 34% | 21 | 26% |
| 2 | 30 | 32% | 79 | 40% | 41 | 51% |
| 3 | 7 | 7% | 16 | 8% | 11 | 14% |
| 4 | 0 | 0% | 5 | 3% | 2 | 2% |
| Total # Animals | 94 | | 197 | | 81 | |

Table S1c Number of hypodermal neoplasias

Experiment 3

| # Growths | Dim | | Unselected | | Bright | |
|---|---|---|---|---|---|---|
| | # Animals | % Animals | # Animals | % Animals | # Animals | % Animals |
| 0 | 42 | 22% | 45 | 18% | 18 | 11% |
| 1 | 62 | 32% | 79 | 32% | 33 | 21% |
| 2 | 68 | 36% | 95 | 38% | 73 | 46% |
| 3 | 19 | 10% | 24 | 10% | 31 | 19% |
| 4 | 0 | 0% | 5 | 2% | 4 | 3% |
| Total # Animals | 191 | | 248 | | 159 | |

### Table S2 Growth to gravid adulthood after 72 hours development on regular NGM

| Experiment | Bright adulthood | Bright total | Bright % adulthood | Dim adulthood | Dim total | Dim % adulthood | Unselected adulthood | Unselected total | Unselected % adulthood |
|---|---|---|---|---|---|---|---|---|---|
| 1 | | | | | | | 37 | 45 | 82% |
| 2 | | | | | | | | | |
| 3 | 46 | 48 | 96% | 46 | 48 | 96% | 48 | 49 | 98% |
| 4 | | | | | | | | | |
| 5 | 38 | 39 | 97% | 38 | 39 | 97% | 36 | 44 | 82% |

### Table S3 Growth to gravid adulthood after 96 hours on high Neomycin concentration NGM

| Experiment | Bright neo adulthood | Bright neo total | Bright neo % adulthood | Dim neo adulthood | Dim neo total | Dim neo % adulthood | Unselected neo adulthood | Unselected neo total | Unselected neo % adulthood |
|---|---|---|---|---|---|---|---|---|---|
| 1 | 15 | 48 | 31% | 2 | 43 | 5% | 1 | 45 | 2% |
| 2 | 26 | 46 | 57% | 1 | 42 | 2% | | | |
| 3 | 25 | 49 | 51% | 15 | 51 | 29% | 9 | 47 | 19% |
| 4 | 25 | 48 | 52% | 15 | 43 | 35% | | | |
| 5 | 6 | 37 | 16% | 2 | 36 | 6% | 1 | 40 | 3% |

**Supplementary Section 4: Persistence of Physiological States.**

**Physiological States are Not Always Persistent.** Our previous reports showed that the high gene expression capacity state revealed by the *hsp-16.2* biomarkers was persistent enough to mean a difference in lifespan[5,8], and also heritable[9]. In the scenario above wherein the animals had a Ras gain of function mutation, animals that developed more neoplasias during L3 larval development expressed more of the *hsp-16.2* biomarker after an adult heat shock. Thus, the state of high Ras expression, or the consequence, was persistent enough to influence adult physiology. Some states may not manifest without some stress and some states may not persist.

We quantified protein expression capacity in embryos and L1 larvae (the time point we measured *eft-3*), and found that the expression level of the *vit-2::GFP* knockin in embryos or $P_{eft\text{-}3}$*::GFP* in those L1 larvae, respectively, did not correlate with adult gene expression capacity. See Supplementary Fig. S16 showing the correlation of L1s and embryos with adult expression levels.

We attempted to determine persistence of other larval states with the adult states. However, we were confounded by current technical and biological limitations. Specifically, we were unable to reliably determine persistence of other larval states (correlation with adult states) because our current anesthesia-based mounting technology confounds our results; we get opposing results with or without anesthesia. And, our no-anesthesia results are currently too technically noisy to rely upon for longitudinal measures with adults for anything other than embryos or L1s.

In our preliminary observations, we were able to determine some L2-L4 larvae are developmentally out of phase, made apparent by the differences in bursting, and some of the in phase larvae have differences in gene expression capacity. How much differences in G or being out of phase (maybe construed as a signaling or timing difference) contribute to subsequent outcomes and establishment of persistent or transient states remains undetermined. So, to reiterate, we observed some animals were at steady state gene expression levels for a given larval phase, and others are still bursting to get into steady larval state; this cell-autonomous bursting was revealed by differential maturation of reporter proteins (animals only deviated in the faster maturing mCherry direction, Supplementary Fig. S17).

In our experiments, the early embryonic/L1 state did not persist, but was consequential (NeoR). The lack of persistence of the embryonic or early L1 state may be due to the fact that embryos can develop to the L1 state without any ribosomes in their genome[10] – suggesting that there is a switch between maternal ribosomes and self-made ribosomes when the animals hatch and exit the L1 diapause and begin feeding. However, the Lehner group's reports support the idea of persistence of physiological states in two ways. The first evidence suggests persistence of physiological states because chaperone reporters could predict subsequent developmental events[11,12]. The second evidence is that differences in the response to larval heat shock persisted into adulthood, affecting ovulation rates[12]. Additionally, we did detect evidence of persistence or some relationship between the larval and adult states when animals bore a Ras gain of function mutation.

**Supplementary Figure S16. High Protein Expression Capacity in the Embryonic or L1 Diapause States Does Not Persist into Adulthood.**

**a.** Average intensity of Peft-3::GFP in the same animals at L1 larval stage and on Day 2 of adulthood is shown. Two panels represent independent experiments. For imaging, animals were mounted on 1% agarose pads and covered with cover slip. No anesthesia was used for L1 larvae; adult animals were anesthetized with 0.2%Tricain/0.02% Tetramisole. L1 larvae were singled onto individual plates after imaging. Panel **b** shows yolk content in 2-cell stage embryos (visualized through knock-in GFP tagged Vit-2 protein) and adult expression of Hsp-90 in the same animals on Day 2 of adulthood. Three panels represent three biological replicates. For imaging, adult animals were anesthetized with 0.2%Tricain/0.02% Tetramisole, mounted on 1% agarose pads and covered with cover slip. Eggs were imaged without anesthesia and then singled onto individual plates. We longitudinally imaged at least twelve animals per experiment.

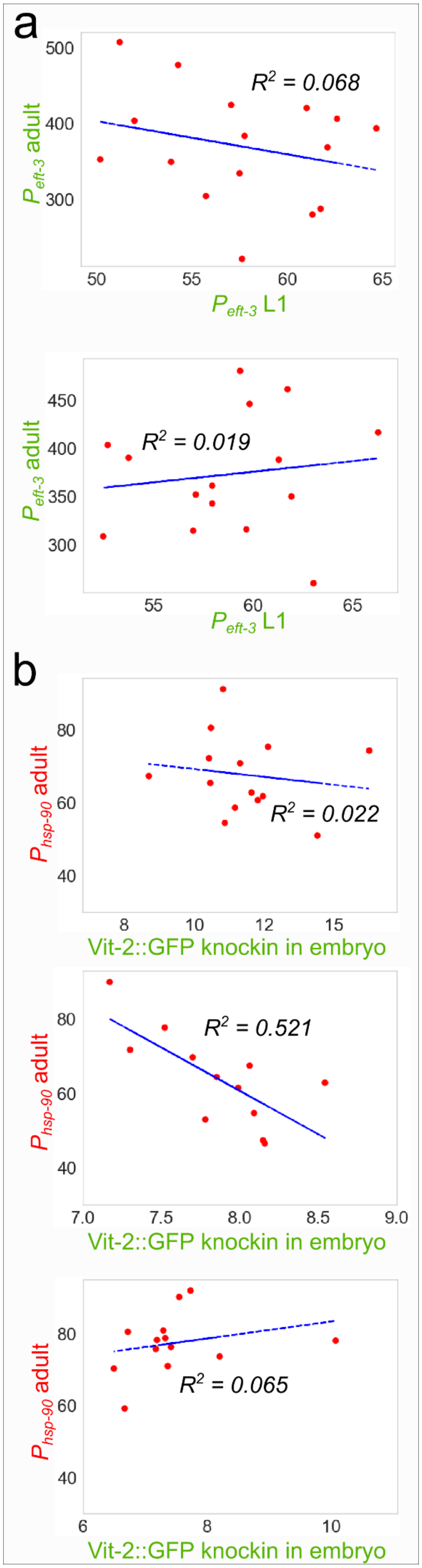

**Supplementary Figure S17. Larval stages exhibit out-of-phase developmental asynchrony punctuated by cell autonomous protein bursting followed by a period of developmentally in phase differences in protein expression capacity.** The left panel of the figure below shows Type I measurements made in young L4 animals where we saw deviation only in the faster maturing mCherry protein, suggesting the animals were not yet in steady state (compare to perfectly yellow animal on the right panel). If we look at animals a few hours later, we see that animals are mostly expressing the same amounts of both alleles, with equivalent minor deviations towards both the mCherry and the mEGFP allele. We saw the same thing with Type II experiments in L2 and L3 animals; deviation only in the mCherry direction in some cells in some animals, followed by a period of seemingly steady state expression wherein the animals varied mostly in the G component. It is currently unclear how much this developmental asynchrony and cell autonomous protein bursting before entry into steady state contribute to differences in biological outcomes. Hence, while our microscopic approach is suitable to investigate gene expression in adults, the same type of analysis in rapidly developing larvae is currently confounded by rapidly changing signaling landscape and bursts of protein synthesis. More technical work will need to be done to quantify these developmental cell autonomous bursting events to determine and their underlying causes and consequences. Specifically, they could be caused by signaling perception differences or expression capacity differences, and they may or may not contribute to or result from physiological states of high or low protein expression capacity or different developmental trajectories. To make these determinations we performed three independent experiments measuring expression in L2 and L4 animals with at least ten animals per group (often dozens). We also performed experiments with L3 animals. In each case, we saw in phase and out of phase animals.

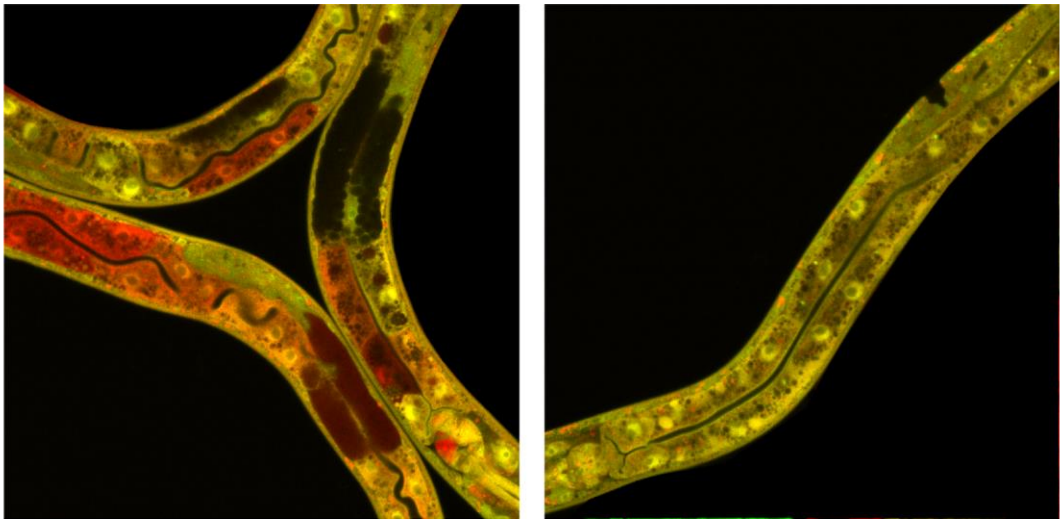

### Supplementary Material Section 5: Trade-offs

**Animals Can Make Trade-Offs, But They May Not Always Do So.** Previous reports showed evidence about trade-offs between stress resistance or lifespan, and progeny production mediated by insulin signaling. One prior reports showed that the insulin signaling system increased lifespan at cost to phenotypic plasticity and progeny production[13,14]. Another report showed that animals that respond better to heat shock as larvae, in terms of lower insulin signaling, have a lowered ovulation rate as adults[12]. So, better response to stress may come at a cost to progeny production. And, we know that the insulin signaling systems also regulates biomarker variation[15] and levels[16], in addition to mediating the adult trade-off between stress resistance/lifespan and progeny production. On the contrary, heat shocks to adults did not reveal trade-offs between animals that responded better or worse (as indicated by the *hsp-16.2* biomarker)[9]. Additionally, no trade-off between self-fertility and lifespan has been observed in unperturbed *C. elegans*[17,18]. One of our working models suggest that animals that express more biomarker do so at the cost of a trade-off with progeny production. Alternatively, animals may express more biomarker simply because the animals that express less are frailer overall. Yet, the dimmer animals did not produce fewer progeny, despite living less time[9]; however, it is unclear if poor responders to heat shock ovulated at the same rate – if they produced more progeny per unit time.

We decided to use an experimental “hammer” to determine if the insulin signaling system could mediate a trade-off after another distinct, but chaperone-related, adult stress. We used genetic models of high and low insulin signaling (*daf-16* and *daf-2,* respectively), and genetic models of lowered nutrient sensing (*nhr-49*) and compromised heat shock response (*hsf-1*); alleles listed in materials and methods. We irradiated wild-type and mutant one day old adult animals with 1000 Joules of UV radiation and measured subsequent lifespan and progeny production. Under these conditions, across genotypes, we saw a tradeoff between progeny production and stress resistance that is mediated by insulin signaling, shown in Supplementary Fig. S18. The figure also shows genetic evidence (the *hsf-1* mutant faring poorly) that the heat shock response was also required for somatic and germline survival; even the seemingly malnourished, sickly *nhr-49* animals did better. Hence, which we were able to observe a trade-off after damaging irradiation with mutants, but not after heat shock within a population of high and low responders.

This data supports a general bet-hedging model wherein insulin signaling mediates interindividual differences in stress perception and/or response resulting trade-offs between somatic maintenance and progeny production among members of isogenic populations, ensuring the fitness of some members of the population in a variety of scenarios. However, it does not negate the fact that we did not detect a trade-off in heat shocked adults in our prior report. It may be that trade-offs cannot be made in wild-type animals after an adult heat shock; that is the notion most supported by the data.

**Supplementary Figure 18. Trade-offs between lifespan and progeny production among different mutants.**

Panel a shows a figure legend. Panels b-d show lifespan after irradiation by 1000 Joules of UV radiation for different *C. elegans* strains. Panels e&f show boxplots of fecundity for five individual animals, with Panel f showing the full dynamic range, including non-irradiated wild-type animals. The boundary of the box closest to zero indicates the 25th percentile, a line within the box marks the median, a dash within the box marks the average, and the boundary of the box farthest from zero indicates the 75th percentile. Whiskers above and below the box indicate the 90th and 10th percentiles. We measured 50 animals in three independent experiments and we measured the fecundity of five individual animals from each genotype.

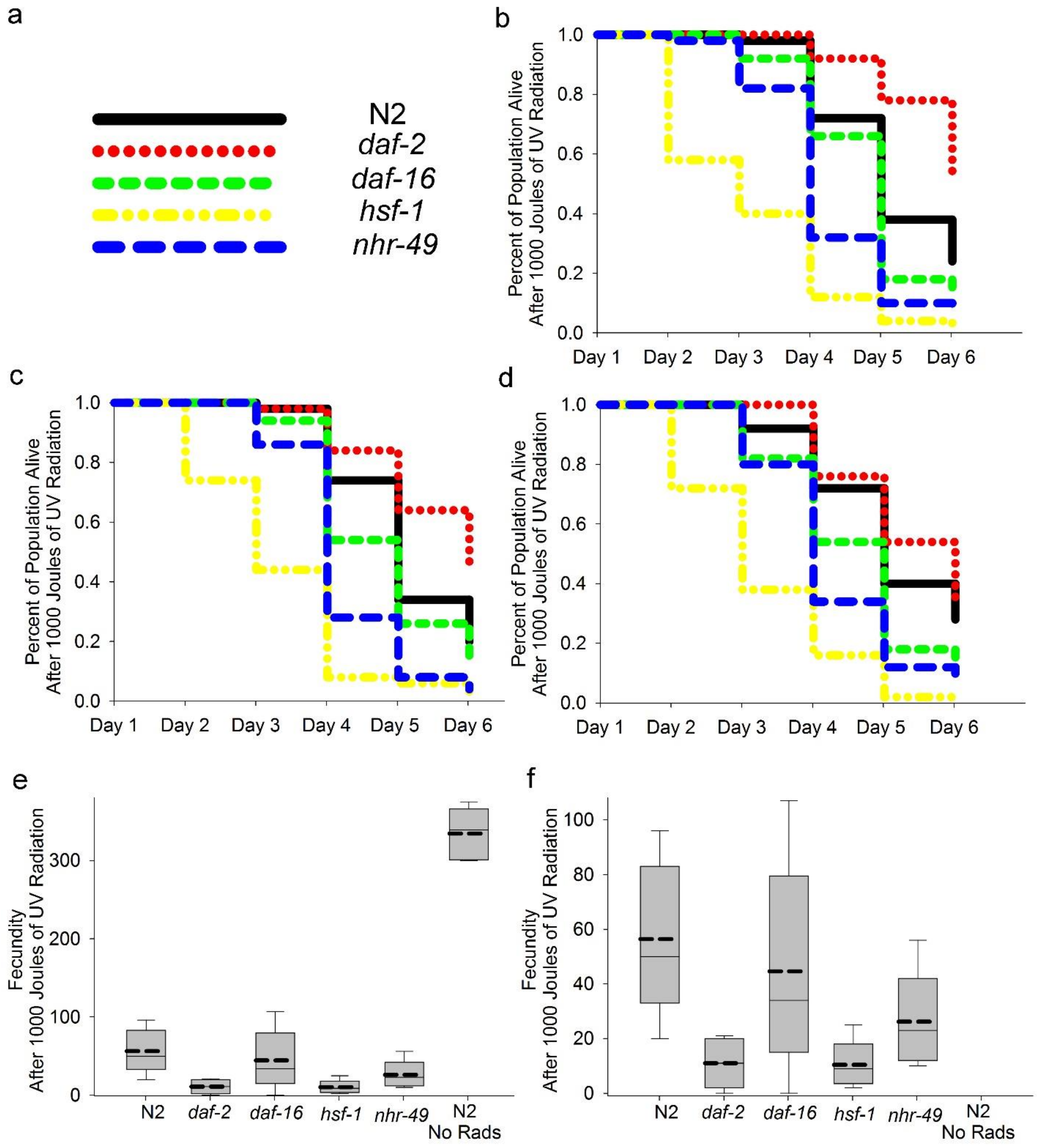
